# Establishment of 3D chromatin structure after fertilization and the metabolic switch at the morula-to-blastocyst transition require CTCF

**DOI:** 10.1101/2021.07.07.451492

**Authors:** Maria Jose Andreu, Alba Alvarez-Franco, Marta Portela, Daniel Gimenez-Llorente, Ana Cuadrado, Claudio Badia-Careaga, Maria Tiana, Ana Losada, Miguel Manzanares

## Abstract

The eukaryotic genome is tightly packed inside the nucleus, where it is organized in 3D at different scales. This structure is driven and maintained by different chromatin states and by architectural factors that bind DNA, such as the multi-zinc finger protein CTCF. Zygotic genome structure is established de novo after fertilization, but the impact of such structure on genome function during the first stages of mammalian development is still unclear. Here, we show that deletion of the Ctcf gene in mouse embryos impairs the correct establishment of chromatin structure, but initial lineage decisions take place and embryos are viable until the late blastocyst stage. Furthermore, we observe that maternal CTCF is not necessary for development. Transcriptomic analyses of mutant embryos show that the changes in metabolic and protein homeostasis programs that occur during the progression from the morula to the blastocyst depend on CTCF. Yet, these changes in gene expression do not correlate with disruption of chromatin structure, but mainly with proximal binding of CTCF to the promoter region of genes downregulated in mutants. Our results show that CTCF regulates both 3D genome organization and transcription during mouse preimplantation development, but mostly as independent processes.

## Introduction

Chromatin is not randomly organized inside the eukaryotic cell nuclei. At the nuclear scale, individual chromosomes occupy distinct spatial territories during interphase (Cremer et al., 2006; Cremer et al., 2010). Within chromosomes, the cell genome is compartmentalized into regions with similar chromatin characteristics, termed ‘A’ and ‘B’ compartments. A-compartments are transcriptionally active, gene-rich and nuclease sensitive; whereas B-compartments are relatively gene poor, transcriptionally silent and insensitive to nucleases (Lieberman-Aiden et al., 2009). At intermediate length scales, high-resolution chromatin interaction maps have revealed that the genome folds into distinct modules called topologically associated domains (TADs). TADs are submegabase sized chromatin domains in which genomic interactions are strong but sharply depleted on crossing the boundary to the adjacent one (Dixon et al., 2012; Hou et al., 2012; Nora et al., 2012; Sexton et al., 2012). Finally, at a smaller scale, distal genomic regions can physically interact forming DNA loop structures that bring distant regulatory elements in close proximity to their target promoters (Dowen et al., 2014; Rao et al., 2014).

During early embryogenesis, there is a massive reorganization of the chromatin. In mature mouse sperm, TADs and compartments are already established (Jung et al., 2017; Du et al., 2017; Ke et al., 2017). However, mature oocytes arrested in metaphase II stage of meiosis are depleted of both structures (Du et al., 2017; Ke et al., 2017), as happens in mitotic cells when chromosomes are highly condensed (Naumova et al., 2013). After fertilization, chromatin adopts a more relaxed state and the strength of TADs and compartments is highly reduced although some weak TADs and interaction domains are detected (Du et al., 2017; Ke et al., 2017; Flyamer et al., 2017; Collombet et al., 2020). During the first cleavage stages, a slow re-establishment of TADs and compartments starts but it is not completed until the 8-cell stage. Moreover, the progressive maturation of chromatin structure is conserved in early animal development, as it has also been observed in *Drosophila* (Hug et al., 2017), zebrafish (Kaaij et al., 2018) and humans (Chen et al., 2019).

A characteristic feature of more than 75% of the borders of TADs and 86% of anchoring points of smaller DNA loops in mammals, is the presence of the CTCF architectural protein (Dixon et al., 2012; Rao et al., 2014; Phillips-Cremins et al., 2013). CTCF is a ubiquitously expressed and highly conserved 11-zinc finger DNA-binding protein (Filippova et al., 1996). It has been shown to be involved in transcriptional regulation, chromatin insulation, genomic imprinting, X-chromosome inactivation, and higher order chromatin organization (Klenova et al., 1993; Bell et al., 1999; Cuddapah et al., 2009; Bell et al., 2000; Hark et al., 2000; Chao et al., 2002; Xu et al., 2007; Dixon et al., 2012). Homozygous CTCF deletion results in early embryonic lethality (Fedoriw et al., 2004; Heath et al., 2008; Wan et al., 2008; Moore et al., 2012; Chen et al., 2019). In somatic cells, conditional knockouts showed additional important roles for CTCF in cell-cycle progression, apoptosis, and differentiation (Heath et al., 2008; Soshnikova et al., 2010; Li et al. 2007; Arzate-Mejia et al., 2018). However, global depletion of CTCF in cell cultures leads to subtle and context-specific changes in gene expression (Zuin et al., 2014; Nora et al., 2017; Hyle et al., 2019; Kubo et al., 2021) although chromatin structure is affected with a reduction of TAD insulation and intra-TAD loop formation. However, we are still missing a clear understanding of the role of CTCF in the initial establishment of chromatin structure, and its impact on regulated gene expression.

Here, we have studied the requirement of CTCF protein during mouse preimplantation development, when 3D chromatin structure is radically reorganized after fertilization, by genetically eliminating the maternal, zygotic, or both contributions of CTCF. To do so, we have performed single-embryo RNA-seq and Hi-C in CTCF mouse mutant blastocysts of different genotypes. Unsuspectedly, maternal mutants are viable and do not show important structural or transcriptional changes compared to control embryos. On the other hand, zygotic and maternal-zygotic mutants die at the late blastocyst stage when they suffer a block in development. Their chromatin structure is affected with an alteration of TAD organization and reduced domain insulation. Transcriptional analysis of maternal-zygotic mutant blastocysts showed a clear downregulation of metabolism related genes and an upregulation of protein homeostasis pathways, but little change to lineage specifiers. Our results show that CTCF, and initial organization of 3D chromatin structure, is not necessary for the first phases of mouse preimplantation development, but after blastocyst cavitation it is absolutely required for developmental progression.

## Results

### CTCF loss leads to embryonic lethality at the late blastocyst stage

In order to analyze the effect of the loss of CTCF during early mouse preimplantation development, we generated both maternal and zygotic mutants. To eliminate maternal contribution of CTCF, we used females carrying a CTCF floxed allele (Heath et al., 2008) and the *Zp3-Cre* driver. This Cre driver is expressed during the growing phase of oocytes prior to the completion of the first meiotic division, and when combined with a floxed allele, it will result in no mRNA or protein generated by the mother to be deposited in the oocyte as it is formed (Lewandoski et al., 1997). Both maternal and maternal-zygotic mutant embryos (M KO and MZ KO) develop up to the blastocyst stage, and they proliferate and undergo cavitation just as zygotic mutant embryos (Z KO) and controls (Fig. 1A). We confirmed the absence of CTCF protein by immunostaining. Z KO present at 8-cell stage a strong reduction of nuclear CTCF and at morula stage CTCF protein is no longer detected in the nuclei (Fig. 1A). On the other hand, M KO embryos exhibit nuclear CTCF protein just as controls, showing that, as they are heterozygous for the deletion, zygotic transcription quickly rescues the lack of maternal CTCF protein. Finally, we confirmed the absence of CTCF protein in MZ KO embryos at all stages examined (Fig. 1A).

**Figure 1.**
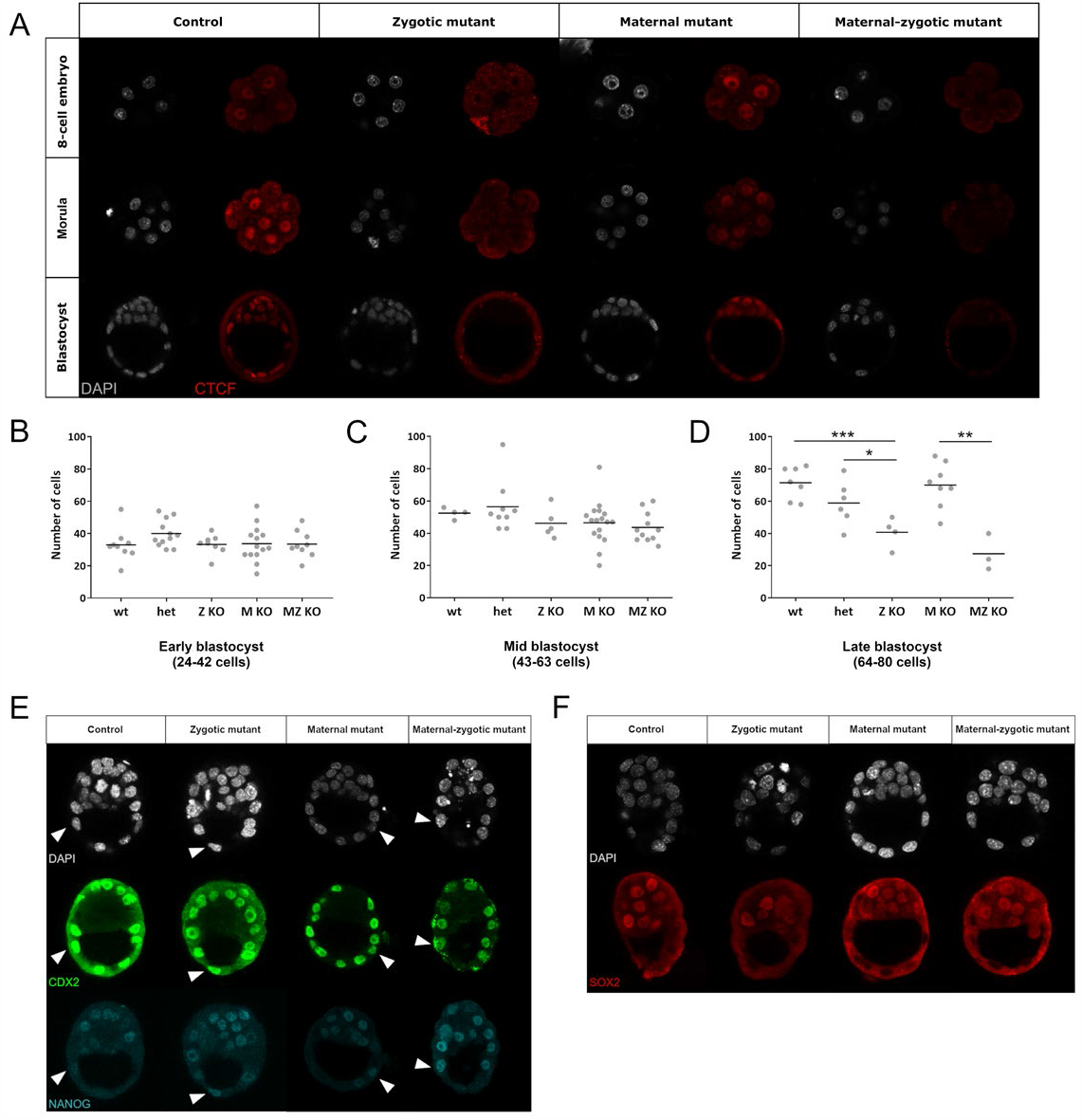
CTCF loss leads to embryonic lethality at the late blastocyst stage. (***A***) Confocal images of control and different combinations of *Ctcf* mutant alleles at 8-cell, morula and blastocyst stages immunostained with CTCF antibody (red). Nuclei were stained with DAPI (gray). (***B****-****D***) Quantification of the total cell number per embryo of early (***B***; 24 to 42 cells), mid (***C***; 43 to 63 cells) and late (***D***; 64 to 80 cells) blastocyst stages for the different *Ctcf* genotypes. Each dot represents an individual embryo. ***p<0.0005, **p<0.005 and *p<0.05 by Student’s t test. (***E***) Confocal images of control (*Ctcf*^*fl/fl*^), zygotic, maternal and maternal-zygotic *Ctcf* mutants immunostained with anti-CDX2 (green) and anti-NANOG (cyan) antibodies. Nuclei were stained with DAPI (gray). Arrowheads indicate cells double positive for CDX2 and NANOG. (***F***) Confocal images of control (*Ctcf*^*fl/fl*^), zygotic, maternal and maternal-zygotic *Ctcf* mutants immunostained with anti-SOX2 antibody (red). Nuclei were stained with DAPI (gray).

Despite being able to form blastocysts, Z and MZ KO embryos suffer a block in development some hours later (Supplemental Fig. S1A). M KO instead develop normally (Supplemental Fig. S1A), are born and grow to adulthood without any apparent phenotype. To determine when CTCF depleted embryos arrest, we subdivided embryos at the blastocyst stage according to cell number and compared them to littermates. We did not detect differences in the number of cells among genotypes at the early-(24-42 cells; Fig. 1B) or mid-blastocyst stage (43-63 cells; Fig. 1C). However, in the late-blastocyst stage (64-80 cells) we observed a striking reduction in the number of cells in Z and MZ KO embryos when compared to their control littermates (Fig. 1D). While Z mutants show a similar median number of cells in late blastocysts as compared to mid-blastocysts (40-45 cells; Fig. 1C, D), MZ mutants have even less cells at this later stage (approximately 30 cells, as compared to 50 at the mid-blastocyst stage; Fig. 1C, D). Next, we examined if this growth arrest is due to a decrease in proliferation or to increased apoptosis. We labeled proliferating cells in mid-blastocyst stage M (with a phenotype equivalent to wild type embryos) or MZ KO embryos with phosphorylated histone H3, and apoptotic cells with TUNEL staining. MZ KO have a tendency to have more apoptotic cells than their littermates (M KO), and we detected a widespread variation in proliferation rates for both genotypes but no significant differences (Supplemental Fig. S1B, C). As CTCF has been described to have a role in maintaining genome stability (Lang et al., 2017), we examined the degree of DNA damage in control and mutant blastocyst. Indeed, we found that there was an increase in γH2Ax staining in nuclear foci in maternal-zygotic mutant as compared to maternal mutants (Supplemental Fig. S1D, E). These results suggest that cells from the preimplantation embryo with no CTCF divide and grow normally, but at certain stage are unable to maintain cell viability, accumulate DNA damage, and are eliminated by apoptosis resulting in embryo lethality.

A possibility that could be occurring was that initial lineage specification in the mouse embryo was not taking place correctly, and that mis-specified cells were leading to embryo arrest. Therefore, we performed immunostainings to detect key factors involved in the commitment of the trophectoderm (CDX2) and the inner cell mass (SOX2 and NANOG) in mid-stage blastocysts. We observed no obvious change in the levels of CDX2 and SOX2 between genotypes, and these were correctly expressed in restricted patterns in the trophectoderm (TE) and the inner cell mass (ICM), respectively (Fig. 1E, F). In contrast, although expression levels remained mostly unchanged, we more frequently observed cells of the TE from Z and MZ KO expressing NANOG (Fig. 1E). We can conclude that although there might be some subtle changes in the expression of NANOG, overall there is no gross mis-specification of the TE and ICM in CTCF depleted blastocysts.

### Reorganization of early chromatin structure in *Ctcf* mutant embryos

To examine the effect of the loss of CTCF on chromatin structure in preimplantation embryos, we developed single blastocyst Hi-C adapting the volumes and timings from single cell Hi-C protocol (Nagano et al. 2015). This was necessary, as at this stage (mid-blastocyst), mutant embryos are phenotypically normal and undistinguishable from littermates (Fig. 1A, C). We therefore needed to generate individual embryo libraries, genotype on the Hi-C sequence data, and then merge individual libraries for the analysis (Supplemental Fig. S2A, B). In order to optimize the number of libraries generated, we used embryos from the cross of *Ctcf* ^*fl/-*^; *Zp3-Cre* ^*tg/+*^ females with *Ctcf* ^*fl/-*^ males, that will produce 50% M KO and 50% of MZ KO (see Methods). As we have shown before that M KO are normal and reach adulthood, we assumed these embryos would be equivalent to wild type controls. Nevertheless, we also included control CD-1 embryos in our study. Overall, we produced 40 high-quality single blastocyst libraries (13 WT, 14 M KO and 13 MZ KO; Supplemental Fig. S2C) with an average of 2.2 million valid contacts each (Supplemental Table S1). Unsupervised clustering of our single blastocyst Hi-C datasets resulted in the classification of MZ KO libraries separately from WT and M KO (Fig. 2A), validating our initial expectation that M KO would be equal to wild type controls. Visual inspection of merged-by-genotype contact probability maps revealed a clear difference in chromatin organization in MZ KO blastocysts when compared to their littermates (M KO) or control embryos (WT), presenting a less defined structure (Fig. 2B). Genome-wide comparison between WT and MZ KO showed a general reduction in insulation, what did not happen when comparing WT to M KO (Fig. 2C). By plotting the frequency of contacts in relation to the distance they span, we observed a noticeable increase in the MZ KO embryos of contacts in the 200 kb to 1 Mb range, that could be associated with TAD organization, as well as a decrease in contacts of 10-50 Mb, a genomic distance associated with compartmentalization (Fig. 2D).

**Figure 2.**
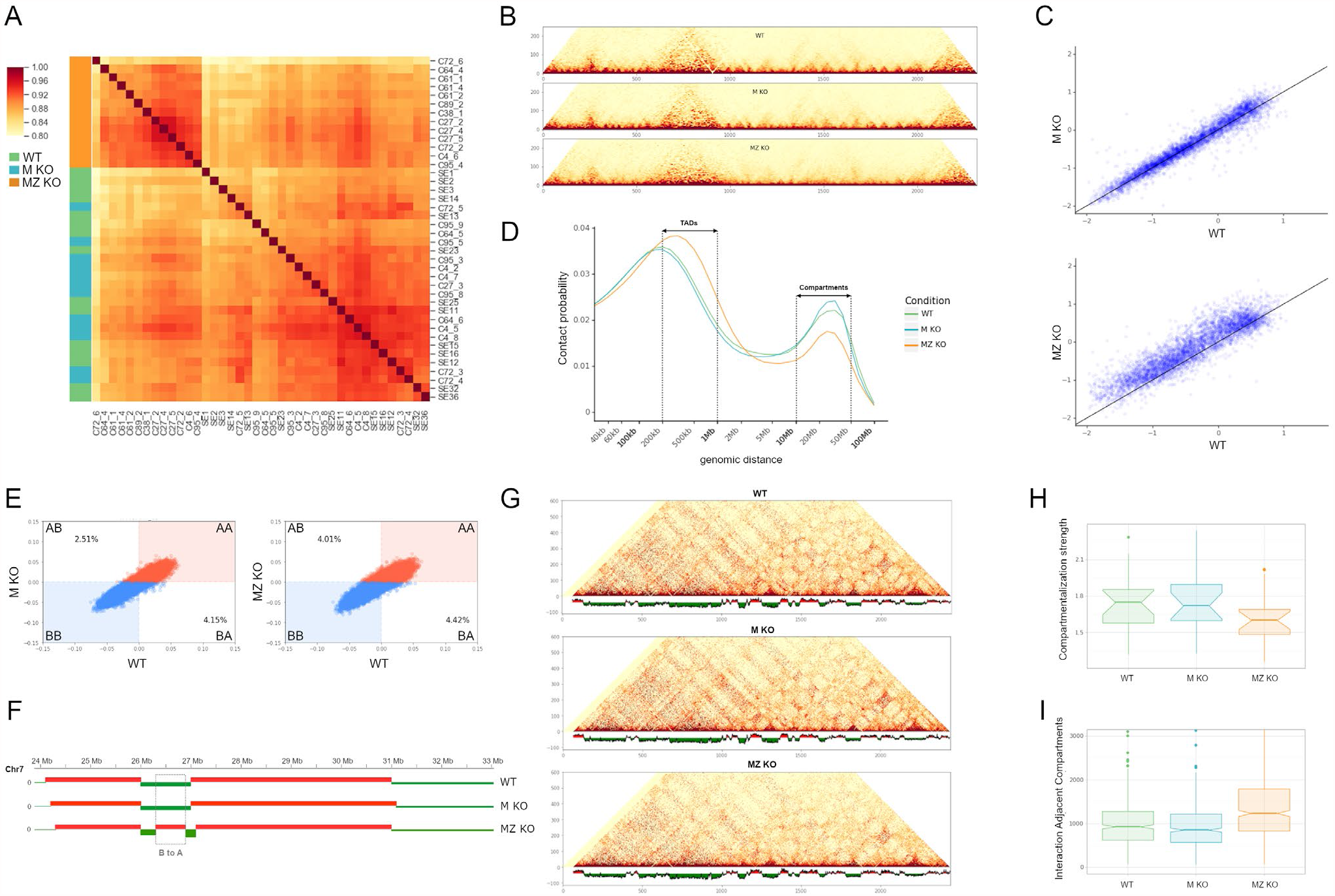
Global reorganization of 3D chromatin structure in *Ctcf* mutant embryos. (***A***) Heatmap of unsupervised clustering of single embryo Hi-C datasets (WT: controls, green; M KO: maternal mutants, blue; MZ KO: maternal-zygotic mutants, orange) by using calculated insulation score with a 500 kb window. (***B***) Coverage-corrected Hi-C matrices of chromosome 17 (chr17:78000000-90000000) from WT, M KO and MZ KO embryos at 40 kb resolution. (***C***) Insulation score values of WT compared to M KO Hi-C datasets (top), and WT compared to MZ KO Hi-C datasets (bottom). Lower insulation scores indicate higher insulation potential. (***D***) Intra-chromosomal contact probability distribution as a function of genomic distance for WT (green line), M KO (blue line) and MZ KO (orange line) embryos. The approximate size of TADs (200 kb t o1 Mb) and compartments (10 Mb to 50 Mb) is indicated. (***E***) Scatterplot of eigenvectors of the intrachromosomal interaction matrices comparing WT and M KO datasets (left) and WT and MZ KO datasets (right). Red represent A compartments and blue B compartments. Numbers within the plot show the percentage of bins that changed compartment from B to A (upper left hand quadrant), or from A to B (lower right hand quadrant). (***F***) Example of a B to A change of compartment in MZ KO embryos in the 26-27 Mb region of chromosome 7. Red and green signals correspond to A and B compartments, respectively. (***G***) Coverage-corrected Hi-C matrices of the full chromosome 11 at 100 kb resolution from WT, M KO and MZ KO embryos showing changes in compartment strength in the MZ KO dataset. The linear arrangement of A (red) and B (green) compartments is shown below each matrix. (***H***) Quantification of changes in compartment strength by calculating the ratios of interaction frequencies between compartments A and B and those between the same compartment for WT, M KO and MZ KO datasets. (***I***) Box plots representing the number of interactions between adjacent compartments in WT, M KO and MZ KO datasets.

Analysis of compartment formation in MZ KO blastocysts revealed that compartments remain mainly unaffected in comparison to WT blastocysts. However, we detected a slight increased frequency of compartments changing their identity from B to A when comparing WT to MZ KO, as opposed to WT compared to M KO (4,01% versus 2,51%; Fig. 2E, F). We also observed an overall reduction in the strength of compartments, as apparent from the less defined checkerboard pattern in MZ KO embryos (Fig. 2G, H), associated with an increased interaction between adjacent compartments in the case of CTCF loss (Fig. 2I).

### TAD assembly in preimplantation development is dependent on CTCF

In addition to the above described changes, MZ KO blastocysts exhibited remarkable differences with control embryos at the TAD organization level. CTCF depleted blastocysts showed a reduction in the number of TADs across the genome leading to an increase in median size (Fig. 3A). We also observe a high degree of TAD reorganization, even in genomic loci that will not be active until later stages in development, such as the *Wnt6-Epha4-Pax3* locus (Supplemental Fig. S3A; Lupiañez et al., 2015), or the HoxD cluster (Supplemental Fig. S3B; Rodriguez-Carballo et al. 2017).

**Figure 3.**
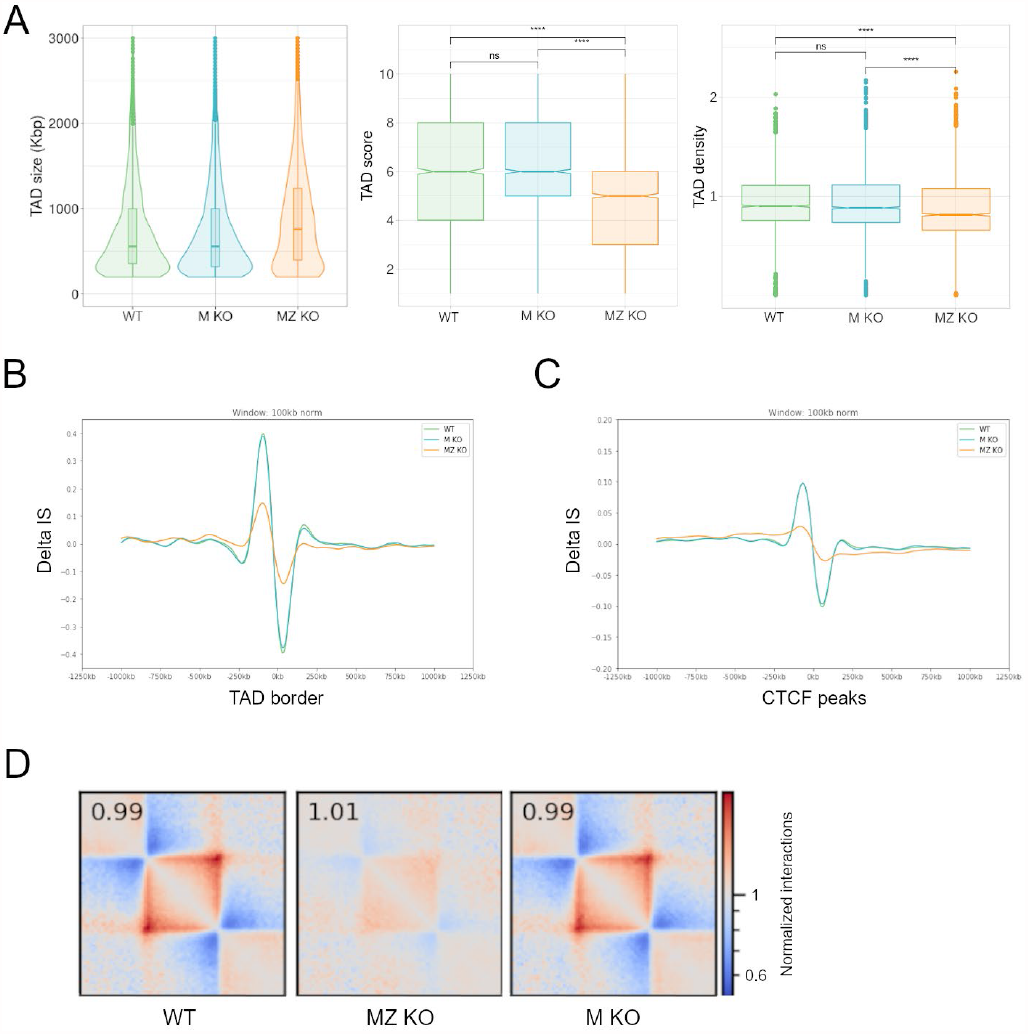
CTCF is required for TAD assembly in blastocysts. (***A***) TAD characterization. Violin and box plots of TAD size (left), TAD reproducibility as score (center) and density of intra-TAD contacts (right). WT (green), M KO (blue), MZ KO (orange). ns, non-significant; ***p<0.0005 by Wilcoxon test. (***B***) Changes in insulation score (Delta IS) at TAD borders in WT (green line), M KO (blue line) and MZ KO (orange line) embryos. Plots are centered on boundary regions ±1000 kb and normalized using a 100 kb window. (***C***) Changes in insulation score (Delta IS) at CTCF peaks in WT (green line), M KO (blue line) and MZ KO (orange line) embryos. Plots are centered on peaks ±1000 kb and normalized using a 100 kb window. (***D***) TAD pile-up contact enrichment map representation in WT, MZ KO and M KO embryos. Each bin size corresponds to 25 kb.

TADs also showed a decrease in their reproducibility score (a statistical measure of how often a TAD is called) and in the density of intra-TAD contacts (Fig. 3A). Changes in insulation score (IS), a measure of the interactions passing across a given genomic position, at the TAD boundaries are less pronounced in MZ KO blastocysts (Fig. 3B) confirming a weaker insulation of these domains when CTCF is not present. Likewise, we detect a reduction in the difference of IS at CTCF peaks when using CTCF ChIP data from naïve embryonic stem (ES) cells (Fig. 3C). Consistent with these results, clear differences in TAD strength are appreciated when data from all TADs are stacked in metaplots, showing a greatly diminished signal in MZ KO as compared to WT of M KO embryos (Fig. 3D).

### Transcriptional changes in blastocyst-stage mutant embryos

To better understand the role of CTCF in preimplantation development and how changes in genomic structure relate to changes in expression, we analyzed the transcriptional changes in CTCF-depleted embryos in single embryos, in order to genotype each embryo on sequence and minimize the loss of starting material. We analyzed mid-blastocyst stage embryos, when mutants for *Ctcf* still do not show any measurable phenotype (Fig. 1A, C), although there is a direct effect on chromatin structure. We studied the expression of control (*Ctcf* ^*fl/-*^ and *Ctcf* ^*fl/fl*^), M KO and MZ KO embryos. Comparison of MZ KO and control embryos revealed approximately 1000 differentially expressed genes (DEG) with 574 genes downregulated and 464 of them upregulated (Fig. 4A; Supplemental Table S2). We did not find any differences when we compared M KO with control embryos that grouped together in principal–component analysis (Supplemental Fig. S5B), in line with our previous observations on the lack of phenotype of the maternal *Ctcf* mutant. KEGG analysis (Kanehisa et al., 2000) of DEG in MZ mutants revealed an enrichment of genes involved in metabolic pathways (glycolysis, such as *Gpi1* or *Slca1*; fatty acid metabolism, *Acadsb* and *Acsl6*; lysosomal enzymes for amino and nucleotide sugar metabolism, *Neu1* and *Hexa*;, oxidative phosphorylation, *Ndufb7, Ndufb10, Ndufab1, Uqcrc1* or *Ppa2*) and DNA replication for downregulated genes; and protein homeostasis (the ribosome, various *Rpl* and *Rps* genes; the proteasome, *Psmd12* or *Sem1*; and the spliceosome, *Hspa* or, *Snrpb*) and cell cycle for upregulated genes (Fig. 4B; Supplemental Table S3). Interestingly, we did not observe changes in lineage determination genes, with the exception of *Nanog*, what matches our previous observations with whole mount immunohistochemistry (Fig. 1E).

**Figure 4.**
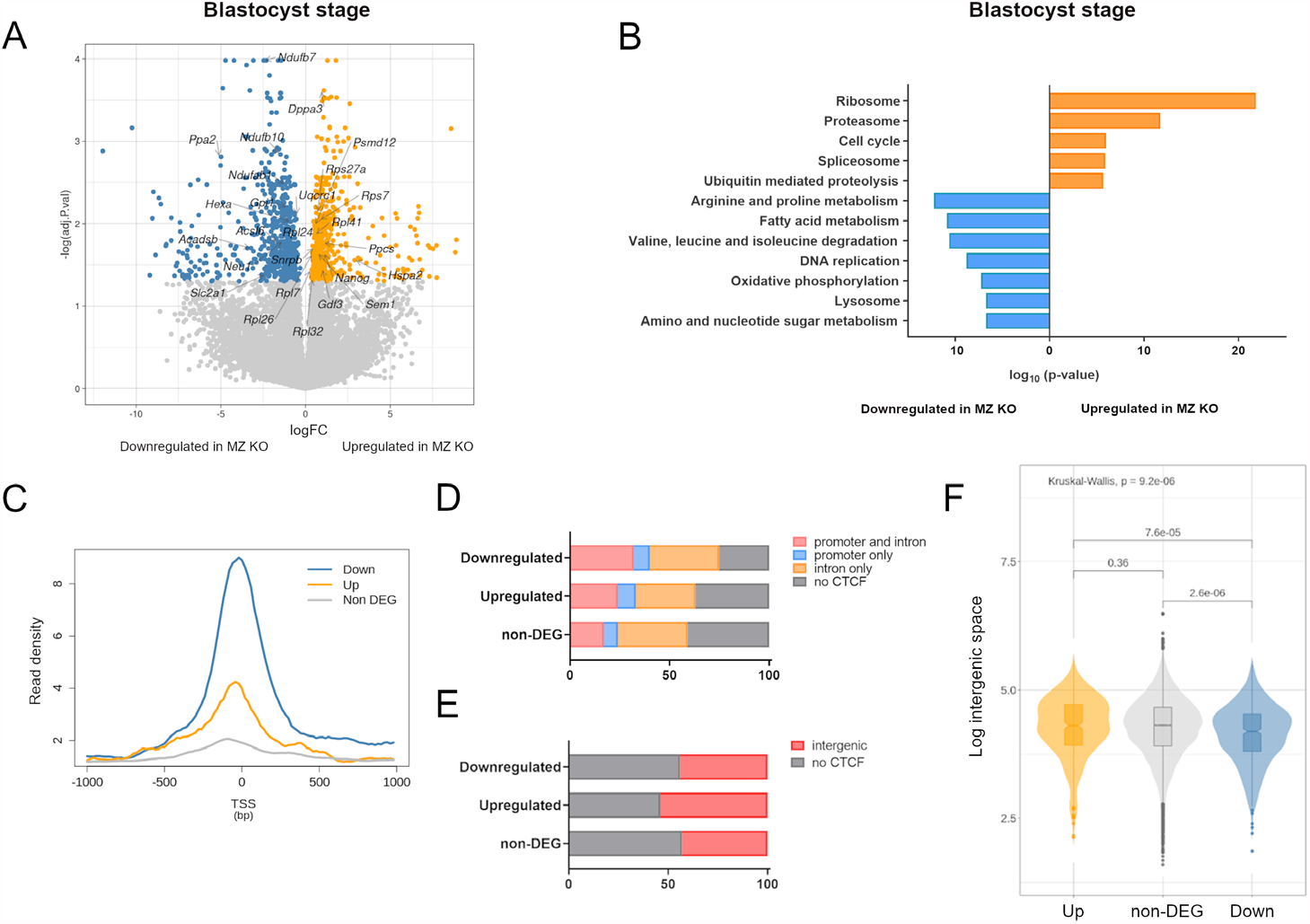
CTCF absence causes transcriptional deregulation at the blastocyst stage. (***A***) Volcano plot of differentially expressed genes between control (*Ctcf*^*fl/fl*^) and MZ KO single blastocysts. Blue indicates downregulated genes in MZ KO (adj. pvalue <0.05 and logFC <−1); orange indicates upregulated genes in MZ KO (adj.pvalue <0.05 and logFC >1) and gray indicates non-differentially expressed genes. Representative genes are indicated. (***B***) Significantly enriched KEGG pathways for genes downregulated (blue) and upregulated (orange) in MZ KO blastocysts. (***C***) Read density plot comparing the ChIP-seq read distribution of CTCF around the TSS (±1000 bp) of downregulated (blue), upregulated (orange) and non-differentially expressed genes (gray) in MZ KO blastocysts. (***D***) Proportion of downregulated, upregulated and non-differentially expressed genes with CTCF binding in the promoter region and an intron (red), only in the promoter (blue) or only in an intron (orange). In gray, genes showing no CTCF binding in promoters or introns. (***E***) Proportion of downregulated, upregulated and non-differentially expressed genes with CTCF binding in intergenic regions (red), or no binding (gray). (***F***) Violin plots showing the distance to the immediately neighboring upstream and downstream genes of downregulated (blue), upregulated (orange), and non-differentially expressed (gray) genes in MZ KO blastocysts. Distances are measured from the transcriptional start or termination sites of each gene, whichever is nearest. Significance calculated by Wilcoxon and Kruskal-Wallis test.

### Promoter binding of CTCF positively regulates gene expression

To characterize the relationship of direct CTCF binding with changes in gene expression, we checked binding of CTCF, as identified by ChIP-seq in naïve ES cells, along the loci of all genes expressed at the blastocyst stage. We first analyzed the distribution of CTCF in a 2 kb region surrounding the transcriptional start site (TSS), finding that downregulated genes have a much higher tendency to have CTCF bound at their promoter region than non-differentially expressed genes. We also observe this trend for upregulated genes, although to a much lesser extent (Fig. 4C). If we extend the search to include introns, we find that downregulated genes are enriched for the co-binding of CTCF to their promoter and at least one intron. In fact, almost 80% of downregulated genes with a CTCF peak on the promoter have also CTCF binding on introns. Strikingly, the proportion of genes bound only at the promoter is similar for up-, downregulated and non-differentially expressed genes in *Ctcf* mutant blastocysts (Fig. 4D). Overall, these results suggest that proximal binding of CTCF to promoters is necessary for positive regulation of gene expression. The observation of co-occurrence or proximal an intron binding is intriguing, and could indicate cooperative mechanisms involving short-range CTCF-mediated interactions.

If we examine intergenic regions, we find an opposite behavior: genes upregulated in mutant blastocyst are more likely to present biding of CTCF in upstream or downstream non-coding regions than downregulated or non-differentially expressed genes (Fig. 4E). This is suggestive of CTCF acting upon cis-regulatory elements located in these regions. However, upregulated genes do not have larger intergenic distances than other expressed genes, as would be expected if these were genes with more complex and numerous intergenic regulatory elements. On the contrary, we did see that downregulated genes tend to be nearer to neighboring genes, and therefore in in more gene-dense regions than non-DEG or upregulated genes (Fig. 4F).

### Reorganization of chromatin structure has a limited impact on transcription

To assess the possible link between structural and transcriptional changes due to the lack of CTCF, we first analyzed changes in the IS surrounding the TSS of DEG. Genes that are downregulated in *Ctcf* mutant blastocysts show a trend for higher insulation at their TSS in normal conditions, as compared to those upregulated or to non-DEG (Fig. 5A). However, these differences can be explained by increased insulation surrounding CTCF bound regions (Fig. 3F), and the more frequent binding of CTCF to promoter regions of DEG, most notably downregulated genes (Fig. 4C). In fact, increased IS is higher in DEG that show binding of CTCF to their promoters (Supplemental Fig. S4A, B). Insulation was diminished in MZ KO, as compared to WT and M KO, for down-and upregulated genes, but did not change in the case of non-DEG (Fig. 5A). Again, this effect was dependent of binding of CTCF to the promoter region, as DEG with no binding hardly changed their IS in the mutants (Supplemental Fig. S3B).

**Figure 5.**
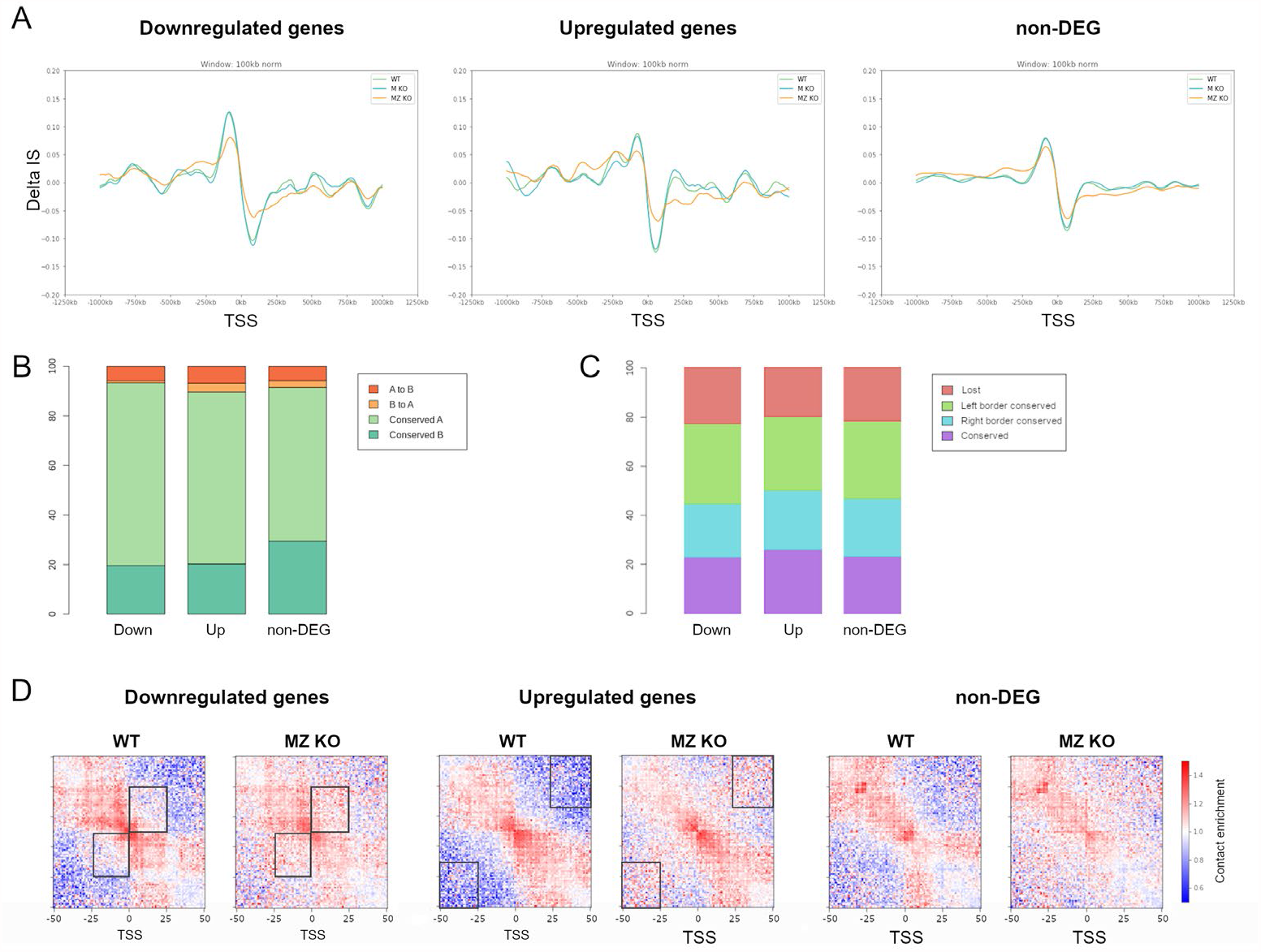
Chromatin reorganization has a limited impact on gene expression. (***A***) Changes in insulation score (Delta IS) at the TSS (±1000 kb) of downregulated (left), upregulated (center) and non-differentially expressed (right) genes in WT (green), M KO (blue) and MZ KO (orange) blastocysts. (***B***) Bar plots showing the percentage of downregulated (DOWN), upregulated (UP), and non-differentially expressed (NON-DEG) genes located in genomic regions where A and B compartments are conserved between controls and MZ KO (light and dark green respectively), or where the identity of the compartment changed from A to B (dark orange) or B to A (light orange). (***C***) Bar plots showing the percentage of downregulated (DOWN), upregulated (UP), and non-differentially expressed (NON-DEG) genes in each of the TAD groups defined by their conservation between WT and MZ KO datasets (conserved, purple; only right boundary conserved, blue; only left boundary conserved, green; not conserved, red). (***D***) Aggregated Hi-C interaction maps centered at the TSS of downregulated (left), upregulated (center) and non-differentially expressed (right) genes that do not show CTCF binding at their promoters in WT and MZ KO embryos. Each bin corresponds to 10 kb.

We next examined the distribution of DEG in compartments, taking into account if they are conserved in MZKO embryos as compared to controls or if they change their identity from A to B and vice versa. We observe that genes that change their expression have more tendency to be localized in A-compartments and less in B-compartments compared to non-DEG (Fig. 5B). Interestingly, downregulated genes are hardly found in compartments that change their identity from B-to-A, as compared to non-DEG or upregulated genes. This could be explained by the fact that changing state from B to A would mean acquiring a permissive chromatin environment for expression. However, we do not observe the opposite behavior, the enrichment of upregulated genes in compartments changing from B to A. On the other hand, DEG and non-DEG are similarly distributed in regions with A-to-B compartment changes (Fig. 5B). We also assigned each expressed gene to a TAD and examined if this was conserved, or one or both boundaries lost, when comparing controls to mutants. In this case, we could not detect differences between DEG and non-DEG genes regarding the behavior of the TAD that contains them upon CTCF loss (Fig. 5C).

Finally, we investigated if changes to local interaction mediated by CTCF binding at regions other than promoters could translate in altered gene expression. To do so, we performed a meta-analysis of the normalized contacts for each genotype centered at the TSS of upregulated, downregulated and non-differentially expressed genes that do not present CTCF binding on the promoter (Fig. 5D). TSS of downregulated genes in MZ KO embryos show a loss of organized short-range interactions (< 250 kb away of the TSS) that likely correspond to enhancer-promoter contacts and could be affecting activation or maintenance of the expression. Instead, TSS of upregulated genes in MZ KO blastocysts exhibit an increase in promiscuous longer-range interactions (> 250 kb away of the TSS). However, we also detect a re-wiring in the interactions at TSS of non-DEG, suggesting that altered expression can partially be explained by changes in local contacts but there are other features that make genes more sensitive to these structural changes.

These results are consistent with what we observe at specific loci. *Slc2a1* gene, encoding the GLUT1 glucose transporter, is downregulated in MZ KO embryos (Fig. 4A), and is localized in a TAD that shows a displacement of one of its borders without affecting the expression of any the immediately neighboring genes. However, the rewiring of contacts at the sub-TAD level results in loss of interactions near *Slc2a1* as can be appreciated in the Hi-C matrix and the Virtual 4C (Supplemental Fig. S4C). The adjacent TAD is split in two in MZ KO blastocysts but only two genes inside of them have its expression affected. For example, *Ppcs*, which encodes an enzyme involved in coenzyme A biosynthesis, is inside one of the split TADs and is the only gene dysregulated inside the TAD, although an increase in contacts is detected in the region (Supplemental Fig. S4C). Similarly, if we focus on the *Nanog* locus, we find some pluripotency genes that together with *Nanog* are upregulated in MZ mutants (*Dppa3, Gdf3*; Fig. 4A, Supplemental Fig. S4D). They are in the same TAD that has its borders expanded and suffers a reshuffling of the local interactions in CTCF depleted embryos (Supplemental Fig. S4D). Curiously, in the same TAD and neighboring *Nanog* sits *Slc2a3*, encoding GLUT3, another glucose transporter, which does not change its transcriptional activity. Together these results suggest that although lack of CTCF affects 3D genome structure at different scales, only local reshuffling of the contacts at the sub-TAD level can partially explain the changes in expression we observe.

### The transcriptional switch of metabolic programs from the morula to the blastocyst is impaired in *Ctcf* mutants

As CTCF has been suggested to have a role in the coordination of dynamic transitions in expression (Sams et al., 2016; Gomez-Velazquez et al., 2017), we investigated if genes changing expression in the morula-to-blastocyst transition were more susceptible to be affected by the lack of CTCF. In first place, we performed single embryo RNA-seq at the early morula stage (16-20 cells; Fig. 1A) of control, Z and MZ mutants. Samples from different genotypes were not resolved by principal– component analysis (Supplemental Fig. S5A), and the comparison of MZ KO with control morulae showed only 55 DEG, most of them being pseudogenes (Supplemental Fig. S5C; Supplemental Table S2). Therefore, we can conclude that the loss of maternal and zygotic CTCF has no effect on transcription before the blastocyst stage, once more reinforcing the notion of its dispensability for the very first stages of mouse development.

We next used our single embryo expression data from controls to examine the transcriptional changes occurring at the morula-to-blastocyst transition. We identified 114 downregulated genes and 702 upregulated genes in this developmental transition (Fig. 6A; Supplemental Table S2). As expected, ICM and embryonic pluripotency genes (such as *Carm1, Dppa1, Tdgf1* or *Zic3*) or TE genes (*Cdx2, Eomes, Gata3*, or *Tfap2α*) (Stirparo et al., 2021), are upregulated from the morula to the blastocyst (Fig. 6A; Supplemental Table S2). However, as observed before (Fig. 1E, F), they do not show much change in *Ctcf* mutant blastocysts (Supplemental Fig. S5D). KEGG analysis showed that several of the pathways and genes changing in the transition to blastocyst are dysregulated in the opposite way to that of MZ KO mutants (Figs 4A, 6A). Oxidative phosphorylation, amino and nucleotide sugar metabolism or lysosome pathways are upregulated from morula to blastocyst (Fig. 6B), while they are were downregulated in MZ KO blastocyst when compared to controls (Fig. 4C). Equally, genes related to the ribosome are downregulated in the blastocyst, but upregulated in *Ctcf* mutants (Figs 4A, 4C, 6A, 6B; Supplemental Table S3).

**Figure 6.**
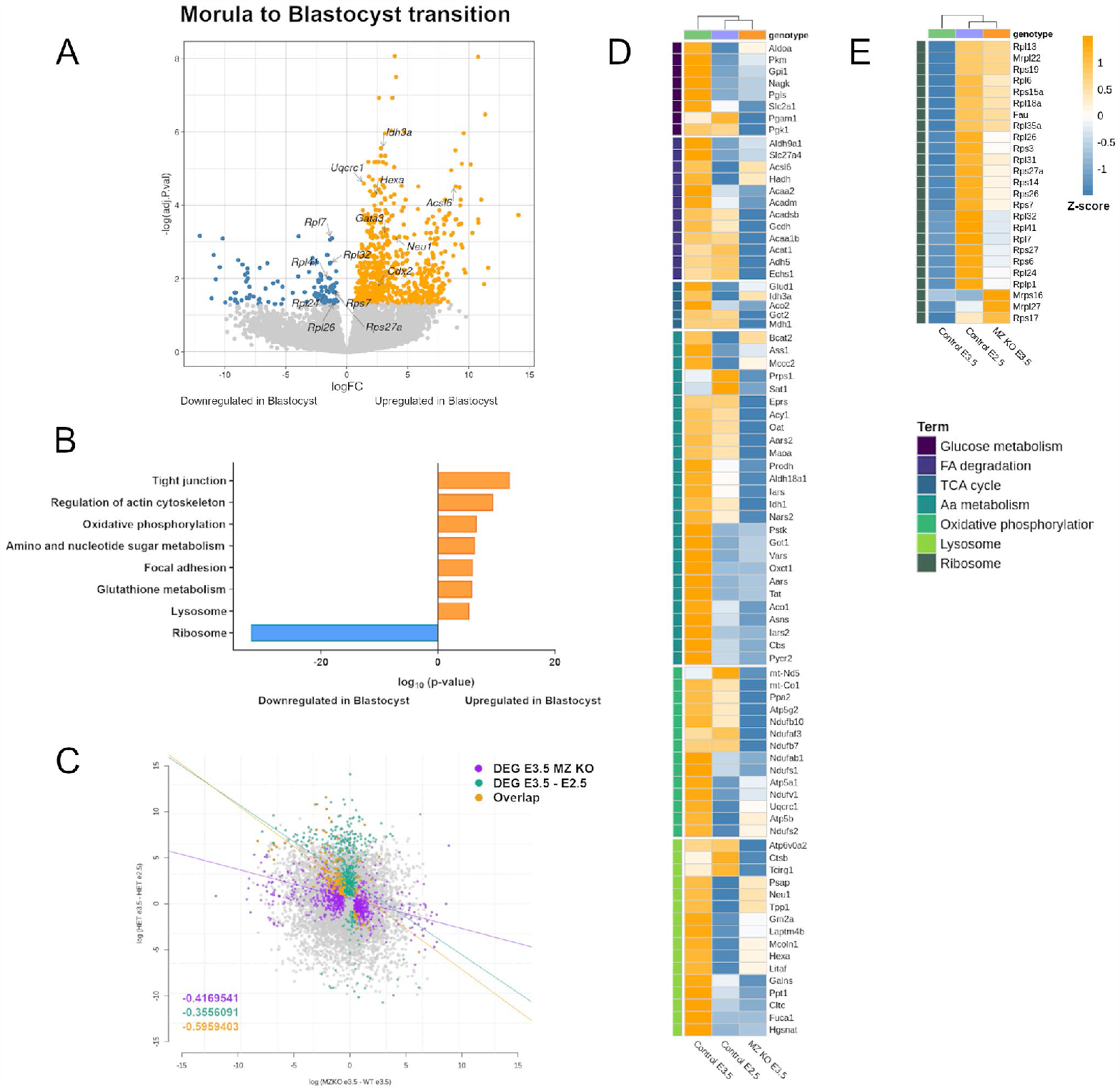
Transcriptional changes occurring from morula to blastocyst are impaired in *Ctcf* mutants. (***A***) Volcano plot of differentially expressed genes between control morulae (*Ctcf*^*fl/-*^) and control blastocysts (*Ctcf*^*fl/-*^). Blue indicates downregulated genes in blastocysts as compared to morulas (adj.pvalue <0.05 and logFC < −1), red indicates upregulated genes in blastocysts as compared to morulas (adj.pvalue <0.05 and logFC >1), and gray indicates non-differential expressed genes. Representative genes are indicated. (***B***) Significantly enriched KEGG pathways for downregulated (blue) and upregulated (orange) genes in control blastocysts as compared to morulas. (***C***) Correlation of the changes in expression between control morulae (*Ctcf*^*fl/-*^) compared to control blastocysts (*Ctcf*^*fl/-*^) in the y-axis, and control blastocysts (*Ctcf*^*fl/fl*^) compared to MZ KO blastocysts in the x-axis. Purple indicates DEG in MZ KO blastocysts only, green indicates DEG in morula-to-blastocyst transition only and orange indicates common DEG for both conditions. Regressions are shown on top of the plot, and correlation values on the bottom left hand corner. (***D***) Heatmap showing the expression (z-scores) of metabolic genes that are downregulated in *Ctcf* mutants, in control morulae (E2.5, purple), control blastocysts (E3.5, green) and MZ KO blastocysts (orange). Clustering of the conditions is shown on top. (***E***) Heatmap showing the expression (z-score) of ribosomal genes that are upregulated in *Ctcf* mutants, in control morulae (E2.5, purple), control blastocysts (E3.5, green) and MZ KO blastocysts (orange). On the right, color code shown on the left side of the heatmaps for KEGG pathways categories. Clustering of the conditions is shown on top.

If we perform a genome-wide comparison of the changes in gene expression of MZ KO blastocysts with the changes that occur during the morula-to-blastocyst transition, we observe a clear inverse correlation for both sets of DEG (Fig. 6C). Moreover, clustering of metabolism and ribosomal genes previously identified as DEG, grouped together E2.5 controls and E3.5 MZKO, with both these showing a similar signature of expression (Fig. 6D, E). Overall, these results suggest that loss of *Ctcf* leads to a failure to correctly implement the changes in gene expression, mainly related to metabolism and protein homeostasis, which would normally occur when the morula progresses to the blastocyst stage.

## Discussion

The role of CTCF in structuring the genome in 3D is undisputed (Ong and Corces, 2014). Equally, the relationship between for chromatin structure and the regulation of gene expression, although subject to debate, is also clear (Ibrahim and Mundlos, 2020). It is therefore surprising the limited transcriptional effect of CTCF deletion, as observed for example in ES cells (Nora et al., 2017; Kubo et al., 2021). This could be the results of CTCF being necessary for the initial establishment of regulatory interactions along chromatin, but not for the posterior maintenance of these contacts. Therefore, depletion of CTCF would not affect established transcriptional programs, but could be needed for changes in gene expression associated to differentiation events (Gomez-Velazquez et al., 2017; Arzate-Mejia et al., 2018). In this view, 3D chromatin structure mediated by CTCF is not required for transcriptional homeostasis, but is essential for cell state transitions that involve deployment of novel expression programs.

We have explored this issue by taking advantage of the first stages of mouse development, starting from fertilization, when the maternal and paternal haploid genomes come together to establish the embryonic diploid genome. We have used mutant embryos with no maternal, zygotic or maternal-zygotic contribution of CTCF, to generate single-embryo Hi-C and RNA-seq data.

We find that maternal CTCF is not required for development or adult life, nor for the establishment of 3D chromatin structure or transcriptional programs in the preimplantation mouse embryo. Our results are in contradiction with previous studies (Wan et al., 2008), surely due to the different approaches used to deplete CTCF (microinjection of RNAi versus genetic deletion). Furthermore, we observe that CTCF is dispensable for the first stages of development, as the block we observe in zygotic and maternal-zygotic mutants only occurs after blastocyst formation. This lack of phenotype suggests that CTCF is not required in the first hours of development before the zygotic gene is expressed and protein produced. Another possibility is that maternal CTCF is rescued by the paternal contribution, as it has been shown that CTCF protein is present on the mature sperm genome, bound to a small number of specific sites (Jung et al., 2017). Nevertheless, this low amount of protein is not sufficient to establish correct chromatin structure in embryos lacking maternal and zygotic CTCF.

Despite limitations in resolution of the contact maps, we observe that complete depletion of CTCF in blastocyst nuclei generates a less organized chromatin structure with an overall reduction in the isolation of interaction domains. Compartment identity remains mainly unaffected, but we have found higher frequency of inter-compartment interactions that translates into a reduction of compartment strength. TADs do not completely disappear in mutant embryos, but again their strength is much reduced and there is a reorganization of domains due to more promiscuous genomic contacts. Indeed, the increase in the number of long-range contacts (200 kb-1 Mb) suggests the formation of larger DNA loops, probably as a consequence of cohesin not encountering obstacles at CTCF binding sites during loop extrusion (Wutz et al., 2020; Hansen 2020).

The fact that interaction domains are established de novo during preimplantation development in the absence of CTCF, argues that other factors or processes must be taking place to initially structure the genome. Similar conclusions have been reached in other systems, as for example the human embryo (Chen et al. 2019). In this scenario, CTCF could be acting as a scaffolding factor that stabilizes and maintains 3D organization generated by other means. Another possibility is that 3D structure is inherited from pre-patterned chromatin present in the separate gamete haploid genomes. However, recent work has shown that the majority of chromatin domains present in the maternal or paternal genomes prior to fertilization are not preserved in the embryo (Collombet et al. 2020). At present, there is no clear candidate/s for this triggering factor of genome structure, and at this point we cannot rule it that it appears as a side consequence of other processes involving genome dynamics, such as DNA replication (Pope et al., 2014) or lamina-chromatin interactions (Borsos et al., 2019).

The absence of CTCF triggers changes in expression related with metabolism and protein homeostasis in the blastocyst, while surprisingly there are no changes yet at the morula stage. Interestingly, in the morula to blastocyst transition there is a metabolic switch in the use of energy (Kaneko, 2016; Zhang et al., 2018) and a downregulation of ribosomal components (Gao et al., 2017). Therefore, changes in the expression of metabolic pathways and structural components of the ribosomes in this transition do not take place in the CTCF mutants, what we believe is the cause of the developmental block of late blastocysts. It is interesting to note that metabolic transitions are important in other developmental transitions in the early mouse embryos, as is the case of the 2C state (Rodriguez-Terrones et al., 2020).

As reported in other contexts where there is a dynamic regulation of expression, lack of CTCF interferes affecting specifically some of the processes involved (Sams et al., 2016; Gomez-Velazquez et al., 2017; Stik et al., 2020). However, it remains unclear by which mechanisms is CTCF facilitating these transcriptional changes.

On the one hand, we have not found a clear relationship between transcriptional changes we observe in mutant embryos and the structural reorganization caused by CTCF absence. Changes in insulation or TAD organization are not translated into global changes of gene expression in those regions, suggesting that transcription is resilient to alterations in TAD domains. Although mutant blastocysts exhibit a rewiring of interactions, these local changes do not alter all the genes in a given region in the same way, indicating that there are other properties that influence the response of genes to modifications of the 3D structure of the genome. The different susceptibility of genes could depend on the proximity between promoters and their enhancers, enhancer-promoter intrinsic preferences (Zabidi et al., 2014; Yokoshi et al., 2020) or if they are genes with dynamic or continuous and stable expression (Sams et al., 2016; Gomez-Velazquez et al., 2017; Stik et al., 2020).

On the other hand, we see a link between CTCF binding at promoters and downregulation of the expression, suggesting another role of CTCF protein different from insulation. In fact, it has been shown that promoter-proximal CTCF can promote long-range enhancer-promoter contacts and facilitate gene activation (Kubo et al., 2021). It is noteworthy that is the case of genes downregulated in mutants, we see enrichment of CTCF bound at introns together with promoters, suggestive of a more complex mechanisms than transcriptional activation by binding to promoter regions.

In summary, we find that CTCF is necessary for a proper 3D organization of the chromatin after fertilization and for transcriptional regulation of the metabolic switch in the morula to blastocyst transition required for developmental progression beyond blastocyst stage. A rather unexpected observation was that we did not see mayor disruption of lineage-specification events or the expression of key factors, such as *Oct4* or *Cdx2*. A notable exception to this was the upregulation of *Nanog*, which occurred together with changes in expression several neighboring genes part of an extended pluripotency cluster (Levasseur et al., 2008). Nevertheless, our mutant embryos, that show an impaired 3D genome structure, were able to properly progress through the first developmental decisions to take place. This argues that the gene regulatory networks responsible for these early events are largely independent of chromatin organization, and must rely on short range regulatory mechanism.

## Materialsand Methods

### Mouse strains

The following mouse lines were used in this work: CD1 (Charles Rivers), Ctcf floxed allele (Heath et al., 2008), Zp3-Cretg/tg (Lewandoski et al., 1997). To obtain controls and zygotic mutant embryos, Ctcf fl/-individuals were inter-crossed. To obtain maternal and maternal-zygotic mutant embryos Zp3-Cre tg/+; Ctcf fl/-females were crossed with Ctcf fl/-males. In this way, all embryos will lack maternal contribution of Ctcf. As for the zygotic Ctcf deletion, 50% will be heterozygous (Ctcf fl/-) and 50% homozygous (Ctcf -/-). Adults were genotyped by PCR of tail-tip DNA using primers detailed in Supplemental Table S4. For preimplantation embryos, genotyping was performed directly on individually isolated embryos after antibody staining.

Mice were bred in the animal facility at the Centro Nacional de Investigaciones Cardiovasculares (Madrid, Spain) in accordance with national and European Legislation.

### Embryo collection and immunofluorescence

Embryos were collected at morula or blastocyst stage by flushing the oviduct or the uterus with M2 medium (Sigma) and fixed for 10 minutes in 4% PFA in PBS. After permeabilization with 0.5% Triton X-100 in PBS (PBST) for 20 minutes, embryos were blocked for 1h with 10% FBS in PBST at room temperature. Incubation with primary antibodies was done overnight at 4ºC in blocking solution and incubation with secondary antibodies was performed for 1h at room temperature in blocking solution. The following antibodies and dilutions were used: mouse monoclonal anti-CTCF (sc-271474, Santa Cruz Biotechnology) 1:200, rabbit monoclonal anti-CTCF (ab128873, Abcam) 1:200, rabbit monoclonal anti-CDX2 (ab76541, Abcam) 1:300, rat monoclonal anti-NANOG (14-5761, eBioscience) 1:200, goat polyclonal anti-SOX2 (AF2018, R&D systems) 1:100, rabbit polyclonal anti-PH3Ser10 (06-570, Millipore) 1:200, mouse monoclonal anti-gamma H2AX (05-636, Millipore) 1:200. Secondary Alexa Fluor conjugated antibodies (Life Technologies) were used at 1:1000. In addition, nuclei were visualized by incubating embryos in DAPI at 1 μg/ml for 15 min.

For TUNEL assay, the Terminal Transferase recombinant kit (Roche 03 333 574 001) and biotin-16-dUTP (Roche 11 093 070 910) were used.

Embryos where imaged on glass-bottomed dishes (Ibidi or MatTek) with a Leica SP5 or Leica SP8 confocal microscope.

### ChIP-seq

4×107 cells growing at 70% of confluence were washed with PBS, trypsinized, resuspended in 20 mL of growing media and cross-linked with 1% formaldehyde for 15 minutes at RT. After quenching with 0.125 M Glycine, fixed cells were washed twice with PBS containing 1 μM PMSF and protease inhibitors, pelleted and lysed in lysis buffer (1%SDS, 10mM EDTA, 50mM Tris-HCl pH 8.1) at 2×107 cells/ml. 107 cells equivalent to 40-50 μg of chromatin were used per immunoprecipitation reaction with 10 μg of antibody. Sonication was performed with a Covaris system (shearing time 20 min, 20% duty cycle, intensity 6, 200 cycles per burst and 30 s per cycle) in a minimum volume of 2 ml. From 6 to 15 ng of immunoprecipitated chromatin (as quantitated by fluorometry) were electrophoresed on an agarose gel and independent sample-specific fractions of 100–200 bp were taken. Adaptor-ligated library was completed by limited-cycle PCR with Illumina PE primers (10-12 cycles). DNA libraries were applied to an Illumina flow cell for cluster generation and sequenced on the Illumina HiSeq2500.

Alignment of sequences to the reference mouse genome (mm10) was performed using ‘ Bowtie2 ′ (version 2.3.3.1) under default settings (Langmead and Salzberg, 2012). Duplicates were removed using Picardtools (version 2.13.2) and peak calling was carried out using MACS2 (version 2.1.1.20160309) after setting the qvalue (FDR) to 0.05 and using the ‘–extsize’ argument with the values obtained in the ‘macs2 predictd’ step (Zhang et al., 2008).

CTCF presence was studied in a 2kb region around the TSS of DEG and non-DEG genes. To avoid the differences in amount of genes between groups, we shuffled 571 genes from the non-DEG group and averaged their signal, repeating the process 1000 times, and finally 571 were randomly selected from this processed list.

Mean read-density profile was performed with deepTools 2.5.4 (Ramírez, et al. deepTools2: a next generation web server for deep-sequencing data analysis, Nucleic Acids Research 2016, 44, W160–W165) and plotted with R v3.6.3 (R Core Team, 2020).

### RNA-seq

RNA-seq was performed on single embryos at the morulae and blastocyst stages. cDNA synthesis was performed using SMART-Seq Ultra Low Input RNA Kit (Clontech). Library preparation and sequencing was performed by the CNIC Genomics Unit using the Illumina HiSeq 2500 sequencer.

Adapters were removed with Cutadapt v1.14 and sequences were mapped with Bowtie2 2.2.9 (Langmead and Salzberg, 2012) and quantified using RSEM v1.3.1 (Li and Dewey, 2011) to the transcriptome set from Mouse Genome Reference GRCm38 and Ensembl Gene Build version 98. We manually add the corresponding genomic sequences of LacZ, Puromycin and Neomycin to our reference genome, fasta and gtf files, for in sillico genotyping. Deferentially expressed genes between groups were normalized and identified using the limma (Ritchie et al., 2015) bioconductor package. We considered as significant, only P-values <0.05 adjusted through Benjamini–Hochberg procedure. To study morula-to-blastocyst transition, RNA-seq libraries of early-(24-42 cells) and mid-blastocyst stage (43-63 cells), were analyzed together by including a common condition in the sequenced libraries an batch as a covariate in the linear model. Molecular Signatures Database (MSigDB) was used for gene enrichment analysis (Liberzon et al., 2011).

### Gene density calculation

Intergenic distances were obtained from Mouse Genome Reference GRCm38 and Ensembl Gene Build version 98. For proper calculation and to avoid artefacts, genes overlapping their neighbors were excluded from the analysis. Pairwise comparison was performed with the use of Wilcoxon test, and multiple groups’ comparison via Kruskal-Wallis test analysis.

### Single embryo Hi-C

We adapted the protocol for single cell Hi-C described in Nagano et al., 2015, optimizing it for single embryo by scaling the reaction volumes and reducing the experimental procedures to avoid sample loss. Briefly, blastocysts were treated with Tyrode acidic solution to remove the zona pellucida, fixed with 2% formaldehyde at room temperature for 10 min and quenched with glycine for 10 min on ice. Blastocysts were then washed with PBS and incubated in 100 μl lysis buffer supplemented with protease inhibitors (10 mM Tris-HCl pH8.0, 10 mM NaCl, 0.2% NP40) on ice for 30 min with occasional mixing. After spinning at 3,000 rpm at 4 °C for 5 min, the pellet was washed with 1X Cutsmart Buffer, gently resuspended in 20 µL of 0.5% SDS in 1 x Cutsmart Buffer and incubated at 65 °C for 10 minutes. SDS was quenched with 40 µL of 1 x Cutsmart Buffer and 12 µL of 10% Triton X-100 at 37 °C for 15 min. Then the chromatin was digested by adding 125 U of MboI and incubated at 37 °C overnight with constant agitation. To label with biotin the digested DNA ends, 0.3 μl of 10 mM dCTP, 0.3 μl of 10 mM dGTP, 0.3 μl of 10 mM dTTP, 7.5 μl of 0.4 mM biotin-14-dATP and 1.6 µL of 5 U/µL DNA Polymerase I Large (Klenow) were added to the solution and the reaction was carried out at 37 °C for 90 min with occasional mixing. Samples were incubated for 15 min at 65 °C to inactivate the Klenow enzyme. After spinning at 3,000 rpm at 4 °C for 5 min, supernatant was discarded and 95.8 μl of water, 12 μl of 10 x NEB T4 DNA ligase buffer, 10 μl 10% Triton X-100, 1.2 μl 10mg/ml BSA and 2000 cohesive end units of T4 DNA ligase (New England Biolabs) were added to ligate the biotin-labelled DNA ends and the reaction was incubated at 16ºC overnight. This was followed by spinning the samples at 3,000 rpm at 4 °C for 5 min and addition of 50 μl of 1 x Cutsmart Buffer and 1 µL of 20 mg/mL Proteinase K for overnight incubation at 65ºC to reverse the cross-link. The biotin-labelled DNA was bound to Dynabeads MyOne Streptavidin T1 (Life Technology) magnetic beads, the washed beads were suspended in 60 µl of 2 x BB buffer (10 mM Tris-HCl pH 7.5, 1 mM EDTA, 2 M NaCl) and added to the single blastocyst sample for 60 min incubation at RT in a rotating wheel. After washing the beads with 1 x BB buffer (5 mM Tris-HCl pH 7.5, 0.5 mM EDTA, 1 M NaCl) and 10 mM Tris-HCl (pH 7.5), the beads were incubated at 37°C with 10 U of Alu I in 1 x Cutsmart Buffer for 60 min with constant agitation for DNA fragmentation. Sequencing library preparation was performed on beads: with 5 U of Klenow Fragment (3’-5’ exo-) in 1 x NEBuffer 2 and 0.2 mM dATP at 37°C for 30 min for the A-tailing reaction, and 800 cohesive end units of T4 DNA ligase (New England Biolabs) in 1 x T4 DNA ligase buffer and 2.5 µL NEBNext adaptor for Illumina (diluted 1/30 from stock) at RT for 60 min, each step with the same washes before and after each reaction. Then the beads were re-suspended in amplification mixture containing 25 µL of water, 25 µL NEBNext High-fidelity PCR Master Mix 2X, 2.5 µL universal primer and 2.5 µL indexed primer and amplified with the program: 95 °C for 2 min, 18 cycles of (95 °C 10 s, 55 °C 30 s, 72 °C 30 s) and 72 °C for 5 min. After amplification, magnetic beads were removed, and the single embryo Hi-C library was purified with Agencourt AMPure XP magnetic beads (Beckman Coulter) and eluted in 12 µl of 10 mM Tris-HCl (pH 8.5). Libraries were sequenced with Illumina HiSeq2500 (50bp, paired end) at the CNAG-CRG Sequencing Unit (Barcelona, Spain), obtaining around 5 million reads each (Supplemental Table S1).

### Single embryo Hi-C data processing and analysis

Data were processed using TADbit (Serra et al., 2017) for read quality control, read mapping, interaction detection, interaction filtering, and matrix normalization. After a FastQC protocol to discard artifacts, the remaining reads were mapped to the reference mouse genome (mm10) using a fragment-based strategy in TADbit. After discarding non-informative contacts -including self-circles, dangling-ends, errors, random breaks or duplicates, those experiments with more than 800.000 valid-pairs were kept. Then, the experiments with the same genotype were merged and normalized. Due to WT and MZ KO merged experiments had similar number of reads, the M KO merged experiment was subsampled for a better comparison among samples. For genotyping on sequence, we considered as MZ KO those embryos whose libraries contained no reads in the 22 kb of the deleted region chr8:105662421-105684451 at Ctcf locus (Heath et al., 2008), using as a control other two random regions of the same size chr8:122710142-122732172 and chr8:122710142-122732172.

TADs were identified by using 40-kb resolution vanilla-normalized and decay-corrected matrices as input to the TAD detection algorithm implemented in TADbit. A–B compartments were detected by calculating the first component of a principal-component analysis (PCA) of chromosome-wide matrices (100kb) and assigning A compartments to the genomic bin with positive PCA1 values and high GC content. Conversely, B compartments were assigned to the genomic bin with negative PCA1 values and low GC content. The insulation score and Delta signal were computed using a custom python script following the same methodology as Crane et al 2015, extracting 10kb resolution raw matrices and using a sliding window of 100kbx100kb. The Observed/Expected metaplots were created using the tool coolpup (Flyamer et al., 2020) and unbalanced matrices. In the case of TAD analysis, 25kb resolution matrices were used with the rescale option to scale all the TADs to the same size. In TSS analysis, the resolution used was 10kb and they are centered in the TSS of the genes. Juicebox software was used for Hi-C data visualization and virtual 4C (Robinson et al., 2016).

### Compartment dynamics and strength

Using 100kb resolution matrices, the number of interactions between A-A, B-B and A-B pairs with greater distance than 2Mb are added. The strength compartmentalization is the result of the following formula (AA+BB)/AB.

Compartment dynamics were studied comparing the identity of each bin of the genome in the MZ KO embryos to its identity in the control embryos. Consecutive regions with the same compartment dynamics were fused, obtaining the coordinates for each block. Overlap between compartment dynamics and DEG and non-DEG genes was performed with bedtools v2.30.0 (Quinlan and Hall, BEDTools: a flexible suite of utilities for comparing genomic features, Bioinformatics 2010, 26, 841–842) and plotted with R v3.6.3 (R Core Team, 2020).

### Statistics

Statistical analysis was performed with an unpaired parametric two-tailed t-test through the software GraphPad Prism v7.04 for Windows (GraphPad Software, USA).

## Supporting information

Tabe S4

## Author Contributions

Conceptualization: MJA, MM; Methodology: MJA, AAF, MP, DGL, AC; Software: AAF, DGL; Validation: MJA, AAF, MP, DGL; Formal Analysis: MJA, AAF, MP, DGL; Investigation: MJA, AAF, MP, DGL, CBC, MT; Data Curation: MJA, AAF, DGL; Writing – Original Draft: MJA, MM; Writing – Review & Editing: MJA, AAF, MP, DGL, AC, CBC, MT, AL, MM; Visualization: MJA, AAF, MP, DGL; Supervision: MJA, AC, AL, MM; Project Administration: MJA, MM; Funding Acquisition: AL, MM.

This article contains supporting information online.

## Acknowledgments

We thank Sneha Gopalan and Tom Fazzio for advice and support; Niels Galjart for the Ctcf mutant mice; and present and past members of the Manzanares lab for support, discussion, and ideas. This work was supported by the Spanish Ministerio de Ciencia e Innovación (grants BFU2017-84914-P and BFU2015-72319-EXP to MM; PID2019-106499RB-I00 to AL). The CBMSO is supported by an Institutional grant from the Fundación Ramon Areces, and the CNIC by the Instituto de Salud Carlos III (ISCIII), the Ministerio de Ciencia e Innovación and the Pro CNIC Foundation.

## References

Arzate-Mejía RG, Recillas-Targa F, Corces VG. 2018. Developing in 3D: the role of CTCF in cell differentiation. Development 145: dev137729.

Bell AC, Felsenfeld G. 2000. Methylation of a CTCF-dependent boundary controls imprinted expression of the Igf2 gene. Nature 405: 482–485.

Bell AC, West AG, Felsenfeld G. 1999. The protein CTCF is required for the enhancer blocking activity of vertebrate insulators. Cell 98: 387–396.

Borsos M, Perricone SM, Schauer T, Pontabry J, de Luca KL, de Vries SS, Ruiz-Morales ER, Torres-Padilla ME, Kind J. 2019. Genome–lamina interactions are established de novo in the early mouse embryo. Nature 569: 729–733.

Chao W, Huynh KD, Spencer RJ, Davidow LS, Lee JT. 2002. CTCF, a candidate trans-acting factor for X-inactivation choice. Science 295: 345–347.

Chen X, Ke Y, Wu K, Zhao H, Sun Y, Gao L, Liu Z, Zhang J, Tao W, Hou Z, et al. 2019. Key role for CTCF in establishing chromatin structure in human embryos. Nature 576: 306–310.

Collombet S, Ranisavljevic N, Nagano T, Varnai C, Shisode T, Leung W, Piolot T, Galupa R, Borensztein M, Servant N, et al. 2020. Parental-to-embryo switch of chromosome organization in early embryogenesis. Nature 580: 142–146.

Cremer T, Cremer M. 2010. Chromosome territories. Cold Spring Harb Perspect Biol 2: a003889.

Cremer T, Cremer M, Dietzel S, Müller S, Solovei I Fakan, S. 2006. Chromosome territories - a functional nuclear landscape. Curr Opin Cell Biol 18: 307–316.

Cuddapah S, Jothi R, Schones DE, Roh TY, Cui K, Zhao K. 2009. Global analysis of the insulator binding protein CTCF in chromatin barrier regions reveals demarcation of active and repressive domains. Genome Res 19: 24–32.

Dixon JR, Selvaraj S, Yue F, Kim A, Li Y, Shen Y, Hu M, Liu JS, Ren B. 2012. Topological domains in mammalian genomes identified by analysis of chromatin interactions. Nature 485: 376–380.

Dowen JM, Fan ZP, Hnisz D, Ren G, Abraham BJ, Zhang LN, Weintraub AS, Schuijers J, Lee TI, Zhao K, et al. 2014. Control of cell identity genes occurs in insulated neighborhoods in mammalian chromosomes. Cell 159: 374–387.

Du Z, Zheng H, Huang B, Ma R, Wu J, Zhang X, He J, Xiang Y, Wang Q, Li Y, et al. 2017. Allelic reprogramming of 3D chromatin architecture during early mammalian development. Nature 547: 232–235.

Du Z, Zheng H, Kawamura YK, Zhang K, Gassler J, Powell S, Xu Q, Lin Z, Xu K, Zhou Q, et al. 2020. Polycomb group proteins regulate chromatin architecture in mouse oocytes and early embryos. Mol Cell 77: 825-839.e7.

Fedoriw AM, Stein P, Svoboda P, Schultz RM, Bartolomei MS. 2004. Transgenic RNAi reveals essential function for CTCF in H19 gene imprinting. Science 303: 238–240.

Filippova GN, Fagerlie S, Klenova EM, Myers C, Dehner Y, Goodwi, G, Neiman PE, Collins SJ, Lobanenkov VV. 1996. An exceptionally conserved transcriptional repressor, CTCF, employs different combinations of zinc fingers to bind diverged promoter sequences of avian and mammalian c-myc oncogenes. Mol Cell Biol 16: 2802–2813.

Flyamer IM, Gassler J, Imakaev M, Brandão HB, Ulianov SV, Abdennur N., Razin SV, Mirny LA, Tachibana-Konwalski K. 2017. Single-nucleus Hi-C reveals unique chromatin reorganization at oocyte-to-zygote transition. Nature 544: 110–114.

Flyamer IM, Illingworth RS, Bickmore WA. 2020. Coolpup.py: versatile pile-up analysis of Hi-C data. Bioinformatics 36: 2980–2985.

Gao Y, Liu X, Tang B, Li C, Kou Z, Li L, Liu W, Wu Y, Kou X, Li J, et al. 2017. Protein expression landscape of mouse embryos during pre-implantation development. Cell Rep 21: 3957–3969.

Gomez-Velazquez M, Badia-Careaga C, Lechuga-Vieco AV, Nieto-Arellano R, Tena JJ, Rollan I, Alvarez A, Torroja C, Caceres EF, Roy AR, et al. 2017. CTCF counter-regulates cardiomyocyte development and maturation programs in the embryonic heart. PLoS Genet 13: e1006985.

Hansen AS. 2020. CTCF as a boundary factor for cohesin-mediated loop extrusion: evidence for a multi-step mechanism. Nucleus 11: 132–148.

Hark AT, Schoenherr CJ, Katz DJ, Ingram RS, Levorse JM, Tilghman SM. 2000. CTCF mediates methylation-sensitive enhancer-blocking activity at the H19/Igf2 locus. Nature 405: 486–489.

Heath H, de Almeida CR, Sleutels F, Dingjan G, van de Nobelen S, Jonkers I, Ling KW, Gribnau J, Renkawitz R, Grosveld F, et al. 2008. CTCF regulates cell cycle progression of αβ T cells in the thymus. EMBO J 27: 2839–2850.

Hou C, Li L, Qin ZS, Corces VG. 2012. Gene density, transcription, and insulators contribute to the partition of the Drosophila genome into physical domains. Mol Cell 48: 471–484.

Houghton FD, Thompson JG, Kennedy CJ, Leese HJ. 1996. Oxygen consumption and energy metabolism of the early mouse embryo. Mol Reprod Dev 44: 476–485.

Hug CB, Grimaldi AG, Kruse K, Vaquerizas JM. 2017. Chromatin architecture emerges during zygotic genome activation independent of transcription. Cell 169: 216-228.e19.

Hyle J, Zhang Y, Wright S, Xu B, Shao Y, Easton J, Tian L, Feng R, Xu P, Li C. 2019. Acute depletion of CTCF directly affects MYC regulation through loss of enhancer-promoter looping. Nucleic Acids Res 47: 6699–6713.

Ibrahim DM, Mundlos S. 2020. The role of 3D chromatin domains in gene regulation: a multi-facetted view on genome organization. Curr Opin Genet Dev 61: 1–8.

Jung YH, Sauria MEG, Lyu X, Cheema MS, Ausio J, Taylor J, Corces VG. 2017. Chromatin states in mouse sperm correlate with embryonic and adult regulatory landscapes. Cell Rep 18: 1366–1382.

Kaaij LJT, van der Weide RH, Ketting RF, de Wit E. 2018. Systemic loss and gain of chromatin architecture throughout zebrafish development. Cell Rep 24: 1-10.e4.

Kanehisa M., Goto S. 2000. KEGG: Kyoto encyclopedia of genes and genomes. Nucleic Acids Res 28: 27–30.

Kaneko KJ. 2016. Metabolism of preimplantation embryo development: a bystander or an active participant? In Current Topics in Developmental Biology, (Academic Press Inc), pp. 259–310.

Ke Y, Xu Y, Chen X, Feng S, Liu Z, Sun Y, Yao X, Li F, Zhu W, Gao L, et al. 2017. 3D chromatin structures of mature gametes and structural reprogramming during mammalian embryogenesis. Cell 170: 367-381.e20.

Klenova EM, Nicolas RH, Paterson HF, Carne AF, Heath CM, Goodwin GH, Neiman PE, Lobanenkov VV. 1993. CTCF, a conserved nuclear factor required for optimal transcriptional activity of the chicken c-myc gene, is an 11-Zn-finger protein differentially expressed in multiple forms. Mol Cell Biol 13: 7612–7624.

Kubo, N, Ishii H, Xiong X, Bianco S, Meitinger, F, Hu R, Hocker JD, Conte M, Gorkin D, Yu M, et al. 2021. Promoter-proximal CTCF binding promotes distal enhancer-dependent gene activation. Nat Struct Mol Biol 28: 152–161.

Lang F, Li X, Zheng W, Li Z, Lu D, Chen G, Gong D, Yang L, Fu J, Shi P Zhou J. 2017. CTCF prevents genomic instability by promoting homologous recombination-directed DNA double-strand break repair. Proc Natl Acad Sci 114: 10912–10917.

Langmead B Salzberg, SL. 2012. Fast gapped-read alignment with Bowtie 2. Nat Methods 9: 357–359.

Leese HJ. 2012. Metabolism of the preimplantation embryo: 40 Years on. Reproduction 143: 417–427.

Levasseur DN, Wang J, Dorschner MO, Stamatoyannopoulos JA, Orkin SH. 2008. Oct4 dependence of chromatin structure within the extended Nanog locus in ES cells. Genes Dev 22: 575–580.

Lewandoski M, Wassarman KM, Martin GR. 1997. Zp3-cre, a transgenic mouse line for the activation or inactivation of IoxP-flanked target genes specifically in the female germ line. Curr Biol 7: 148–151.

Li B, Dewey CN. 2011. RSEM: accurate transcript quantification from RNA-Seq data with or without a reference genome. BMC Bioinformatics 12: 323.

Li T, Lu L. 2007. Functional role of CCCTC binding factor (CTCF) in stress-induced apoptosis. Exp Cell Res 313: 3057–3065.

Liberzon A, Subramanian A, Pinchback R, Thorvaldsdottir H, Tamayo P, Mesirov JP. 2011. Molecular signatures database (MSigDB) 3.0. Bioinformatics 27: 1739–1740.

Lieberman-Aiden E, Van Berkum NL, Williams L, Imakaev M, Ragoczy T, Telling A, Amit I, Lajoie BR, Sabo PJ, Dorschner MO, et al. 2009. Comprehensive mapping of long-range interactions reveals folding principles of the human genome. Science 326: 289–293.

Lupiáñez DG, Kraft K, Heinrich V, Krawitz P, Brancati F, Klopocki E, Horn D, Kayserili H, Opitz JM, et al. 2015. Disruptions of topological chromatin domains cause pathogenic rewiring of gene-enhancer interactions. Cell 161: 1012–1025.

Moore JM, Rabaia NA, Smith LE, Fagerlie S, Gurley K, Loukinov D, Disteche CM, Collins SJ, Kemp CJ, Lobanenkov VV, et al. 2012. Loss of maternal CTCF is associated with peri-implantation lethality of Ctcf null embryos. PLoS One 7: e34915.

Nagano T, Lubling Y, Yaffe E, Wingett SW, Dean W, Tanay A, Fraser P. 2015. Single-cell Hi-C for genome-wide detection of chromatin interactions that occur simultaneously in a single cell. Nat Protoc 10: 1986–2003.

Naumova N, Imakaev M, Fudenberg G, Zhan Y, Lajoie BR, Mirny LA, Dekke, J. 2013. Organization of the mitotic chromosome. Science 342: 948–953.

Nora EP, Goloborodko A, Valton AL, Gibcus JH, Uebersohn A, Abdennur N, Dekker J, Mirny LA, Bruneau BG. 2017. Targeted degradation of CTCF decouples local insulation of chromosome domains from genomic compartmentalization. Cell 169: 930-944.e22.

Nora EP, Lajoie BR, Schulz EG, Giorgetti L, Okamoto I, Servant N, Piolot T, Van Berkum NL, Meisig J, Sedat J, et al. 2012. Spatial partitioning of the regulatory landscape of the X-inactivation centre. Nature 485: 381–385.

Ong CT, Corces VG. 2014. CTCF: An architectural protein bridging genome topology and function. Nat Rev Genet 15: 234–246.

Phillips-Cremins JE, Sauria MEG, Sanyal A, Gerasimova TI, Lajoie BR, Bell JSK, Ong CT, Hookway TA, Guo C, Sun Y, et al. 2013. Architectural protein subclasses shape 3D organization of genomes during lineage commitment. Cell 153: 1281–1295.

Pope BD, Ryba T, Dileep V, Yue F, Wu W, Denas O, Vera DL, Wang Y, Hansen RS, Canfield TK, et al. 2014. Topologically associating domains are stable units of replication-timing regulation. Nature 515: 402–405.

Quinlan AR, Hall IM. 2010. BEDTools: a flexible suite of utilities for comparing genomic features. Bioinformatics 26: 841–842.

Rao SSP, Huntley MH, Durand NC, Stamenova EK, Bochkov ID, Robinson JT, Sanborn AL, Machol I, Omer AD, Lander ES, et al. 2014. A 3D map of the human genome at kilobase resolution reveals principles of chromatin looping. Cell 159: 1665–1680.

Ritchie ME, Phipson B, Wu D, Hu Y, Law CW, Shi W, Smyth GK. 2015. limma powers differential expression analyses for RNA-sequencing and microarray studies. sNucleic Acids Res 43, e47.

Robinson JT, Turner D, Durand NC, Thorvaldsdóttir H, Mesirov JP, Aiden EL. 2018. Juicebox.js provides a cloud-based visualization system for Hi-C data. Cell Syst 6: 256-258.e1.

Rodríguez-Carballo E, Lopez-Delisle L, Zhan Y, Fabre PJ, Beccari L, El-Idrissi I, Huynh THN, Ozadam H, Dekker J, Duboule D. 2017. The HoxD cluster is a dynamic and resilient TAD boundary controlling the segregation of antagonistic regulatory landscapes. Genes Dev 31: 2264–2281.

Rodriguez-Terrones D, Hartleben G, Gaume X, Eid A, Guthmann M, Iturbide A, Torres-Padilla ME. 2020. A distinct metabolic state arises during the emergence of 2-cell-like cells. EMBO Rep 21: e48354.

Sams DS, Nardone S, Getselter D, Raz D, Tal M, Rayi PR, Kaphzan H, Hakim O, Elliott E. 2016. Neuronal CTCF is necessary for basal and experience-dependent gene regulation, memory formation, and genomic structure of BDNF and Arc. Cell Rep 17: 2418–2430.

Serra F, Baù D, Goodstadt M, Castillo D, Filion GJ, Marti-Renom MA. 2017. Automatic analysis and 3D-modelling of Hi-C data using TADbit reveals structural features of the fly chromatin colors. PLoS Comput Biol 13: e1005665.

Sexton T, Yaffe E, Kenigsberg E, Bantignies F, Leblanc B, Hoichman M, Parrinello H, Tanay A, Cavalli G. 2012. Three-dimensional folding and functional organization principles of the Drosophila genome. Cell 148: 458– 472.

Soshnikova N, Montavon T, Leleu M, Galjart N, Duboule D. 2010. Functional analysis of CTCF during mammalian limb development. Dev Cell 19: 819– 830.

Splinter E, Heath H, Kooren J, Palstra RJ, Klous P, Grosveld F, Galjart N, de Laat W. 2006. CTCF mediates long-range chromatin looping and local histone modification in the β-globin locus. Genes Dev 20: 2349–2354.

Stik G, Vidal E, Barrero M, Cuartero S, Vila-Casadesús M, Mendieta-Esteban J, Tian TV, Choi J, Berenguer C, Abad A, et al. 2020. CTCF is dispensable for immune cell transdifferentiation but facilitates an acute inflammatory response. Nat Genet 52: 655–661.

Stirparo GG, Kurowski A, Yanagida A, Bates LE, Strawbridge SE, Hladkou S, Stuart HT, Boroviak TE, Silva JCR, Nichols J. 2021. OCT4 induces embryonic pluripotency via STAT3 signaling and metabolic mechanisms. Proc Natl Acad Sci 118: e2008890118.

Wan LB, Pan H, Hannenhalli S, Cheng Y, Ma J, Fedoriw A, Lobanenkov V, Latham KE, Schultz RM, Bartolomei MS. 2008. Maternal depletion of CTCF reveals multiple functions during oocyte and preimplantation embryo development. Development 135: 2729–2738.

Wutz G, Ladurner R, St Hilaire BG, Stocsits RR, Nagasaka K, Pignard B, Sanborn A, Tang W, Varnai C, Ivanov MP, et al. 2020. ESCO1 and CTCF enable formation of long chromatin loops by protecting cohesinstag1 from WAPL. eLife 9: e52091.

Xu N, Donohoe ME, Silva SS, Lee JT. 2007. Evidence that homologous X-chromosome pairing requires transcription and Ctcf protein. Nat Genet 39: 1390–1396.

Yokoshi M, Segawa K, Fukaya T. 2020. Visualizing the role of boundary elements in enhancer-promoter communication. Mol Cell 78: 224-235.e5.

Zabidi MA, Arnold CD, Schernhuber K, Pagani M, Rath M, Frank O, Stark A. 2015. Enhancer-core-promoter specificity separates developmental and housekeeping gene regulation. Nature 518: 556–559.

Zhang J, Zhao J, Dahan P, Lu V, Zhang, C, Li H, Teitell MA. 2018. Metabolism in pluripotent stem cells and early mammalian development. Cell Metab 27: 332–338.

Zuin J, Dixon JR, van der Reijden MI, Ye Z, Kolovos P, Brouwer RW, van de Corput, MP, van de Werken HJ, Knoch TA, van Ijcken, WF, et al. 2014. Cohesin and CTCF differentially affect chromatin architecture and gene expression in human cells. Proc Natl Acad Sci 111: 996–1001.

