## Supplementary Material for "Establishment of 3D chromatin structure after fertilization and the metabolic switch at the morula-to-blastocyst transition require CTCF"

**Andreu et al.**

### Supplementary Figures S1-S5

### Supplementary Tables S1-S4

**Table S1.** Total read and valid pairs of individual single-embryo Hi-C libraries used in this study.

**Table S2.** Differentially expressed genes in *Ctcf* mutants and between wild-type morula and blastocyst stage embryos.

**Table S3.** KEGG pathway enrichment in differentially expressed genes in *Ctcf* mutants and between wild-type morula and blastocyst stage embryos.

**Table S4.** Primers used for genotyping of embryos.

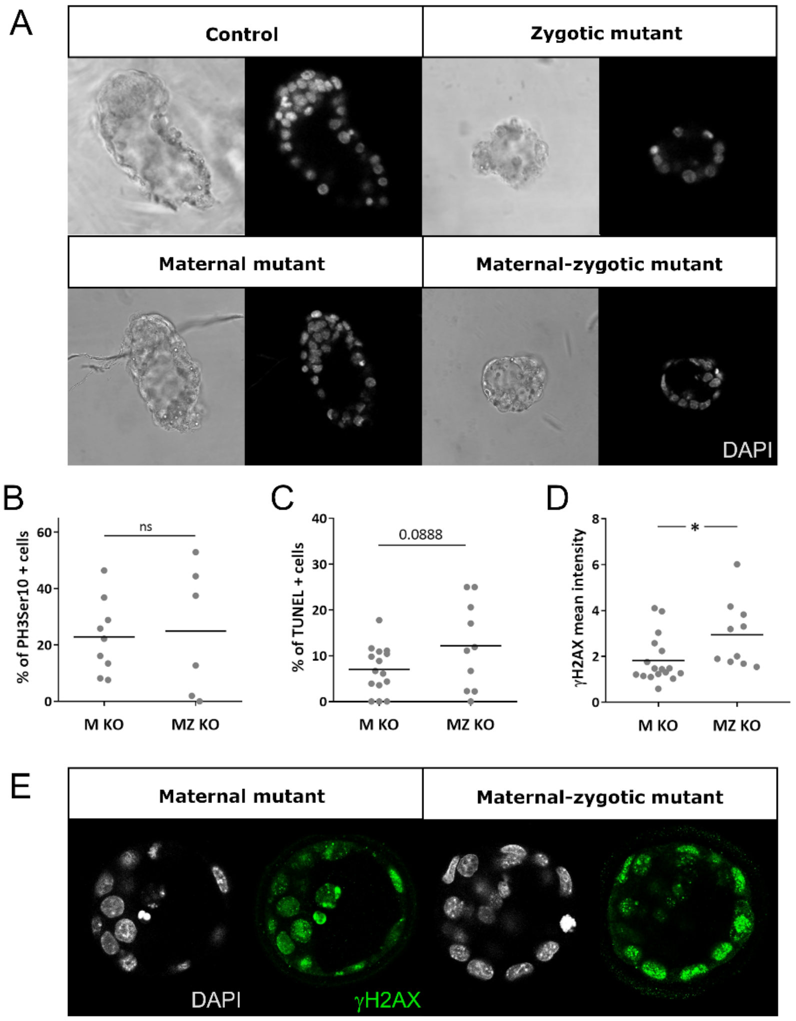


**Figure S1. Phenotype of *Ctcf* mutants.** (***A***) Bright-field and confocal images of late blastocyst control (*Ctcf*$^{fl/-}$), zygotic, maternal and maternal-zygotic *Ctcf* mutants embryos. Nuclei were stained with DAPI. (***B***) Scatter plot of percentage of PH3Ser10 positive cells per embryo in M KO (n=10) and MZ KO (n=6) blastocysts. ns, not significant by Student's t test. (***C***) Scatter plot of percentage of TUNEL positive cells per embryo in M KO (n=15) and MZ KO (n=10) blastocysts. p=0.0888 by Student's t test. (***D***) Scatter plot of anti-γH2AZ signal in in M KO (n=16) and MZ KO (n=10) blastocysts. *, p>0.05 Student's t test. (*E*) Confocal images of maternal and maternal-zygotic mutant embryos stained with an anti-γH2AZ antibody to detect DNA damage (green). Nuclei were stained with DAPI (gray).

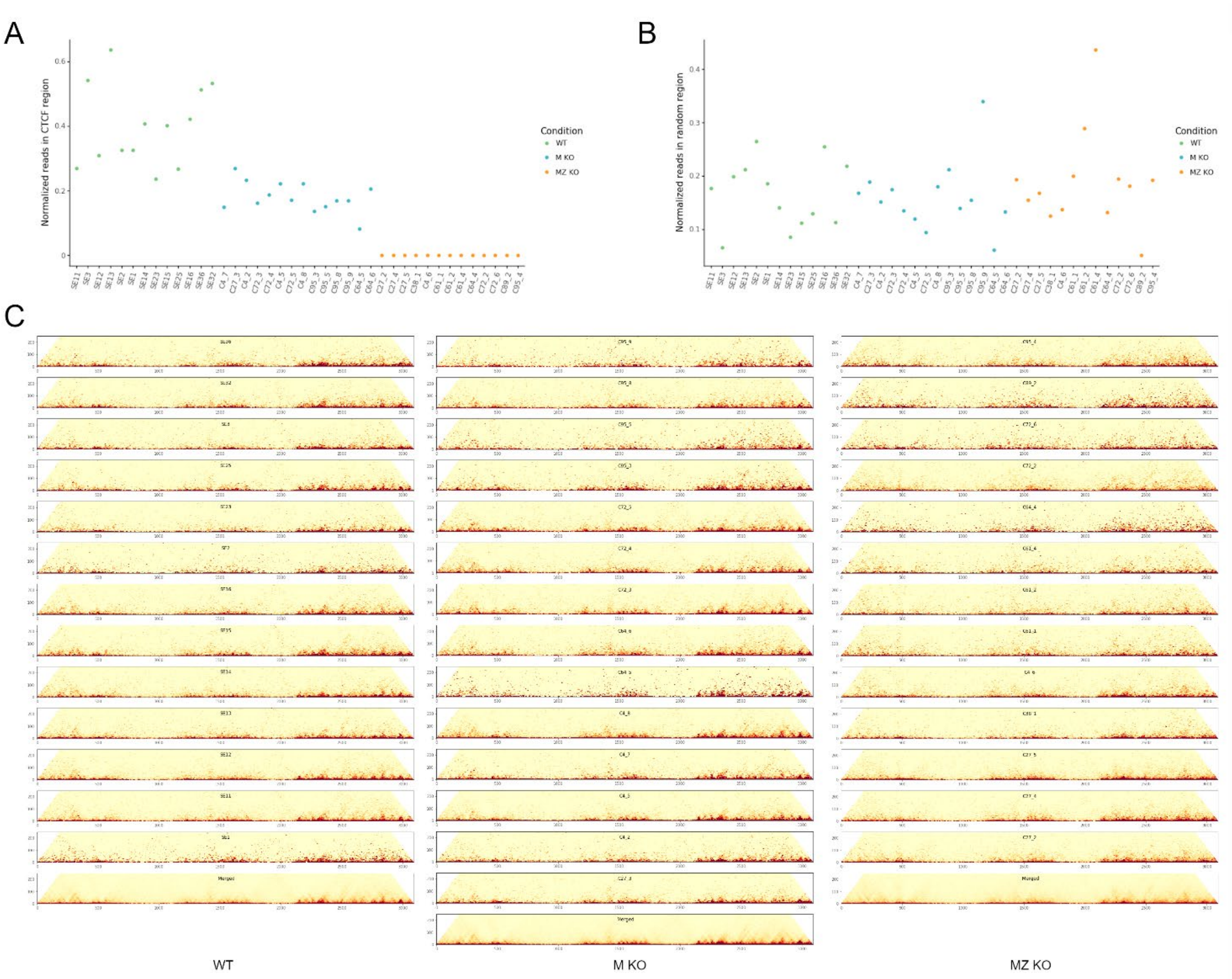


**Figure S2. Single blastocyst Hi-C libraries of WT, M KO and MZ KO embryos.** (*A*) Normalized number of reads at *Ctcf* genomic region (y-axis) for all single blastocyst Hi-C libraries (x-axis), calculated as the number of reads of the deleted region (chr8:105662421-105684451) relative to the number of reads of a region of the same size (chr8:122710142-122732172). WT embryos are shown in green, M KO embryos in blue and MZ KO embryos in orange. (***B***) Normalized number of reads at a random genomic region (y-axis) for all single blastocyst Hi-C libraries (x-axis), calculated as the number of reads of the random region (chr8:75657421-75679451) relative to the number of reads of a region of the same size (chr8:122710142-122732172). WT embryos are shown in green, M KO embryos in blue and MZ KO embryos in orange. (***C***) Single embryo and merged (bottom) Hi-C matrices of chromosome 17 at 40 kb resolution from each individual WT (left), M KO (center) and MZ KO (right) blastocyst.

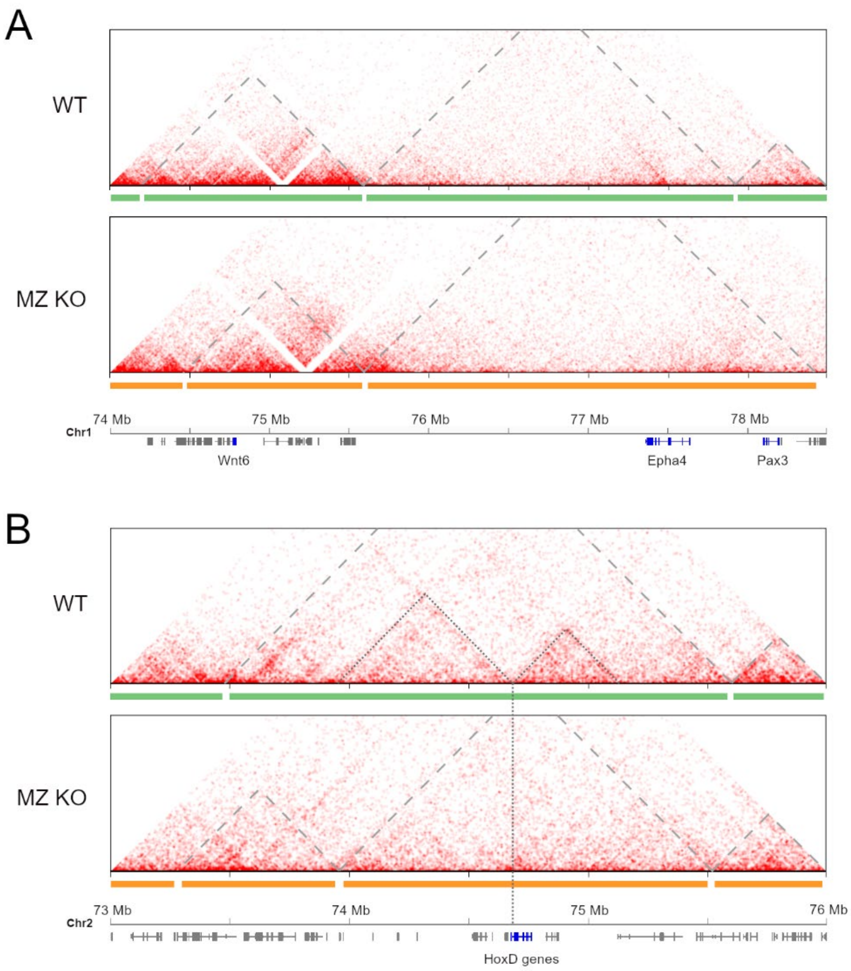


**Figure S3. TAD reorganization of genomic loci not active in the preimplantation embryo.** (***A***) Hi-C contact maps of the *Wnt6-Epha4-Pax3* locus from WT and MZ KO blastocysts. TAD domains in WT and MZ KO embryos are indicated below the matrices in green and orange, respectively. *Wnt6*, *Epha4* and *Pax3* genes are highlighted in dark blue on the gene map shown at the bottom. Dashed lines in gray highlight TAD domains in the matrices. (***B***) Hi-C contact maps of the *HoxD* locus from WT and MZ KO blastocysts. TAD domains in WT and MZ KO embryos are indicated below the matrices in green and orange, respectively. The *HoxD* cluster is highlighted in blue on the gene map shown at the bottom. Dashed lines in gray highlight TAD domains in the matrices and dotted lines show sub-domains inside the TAD containing the *HoxD* cluster.

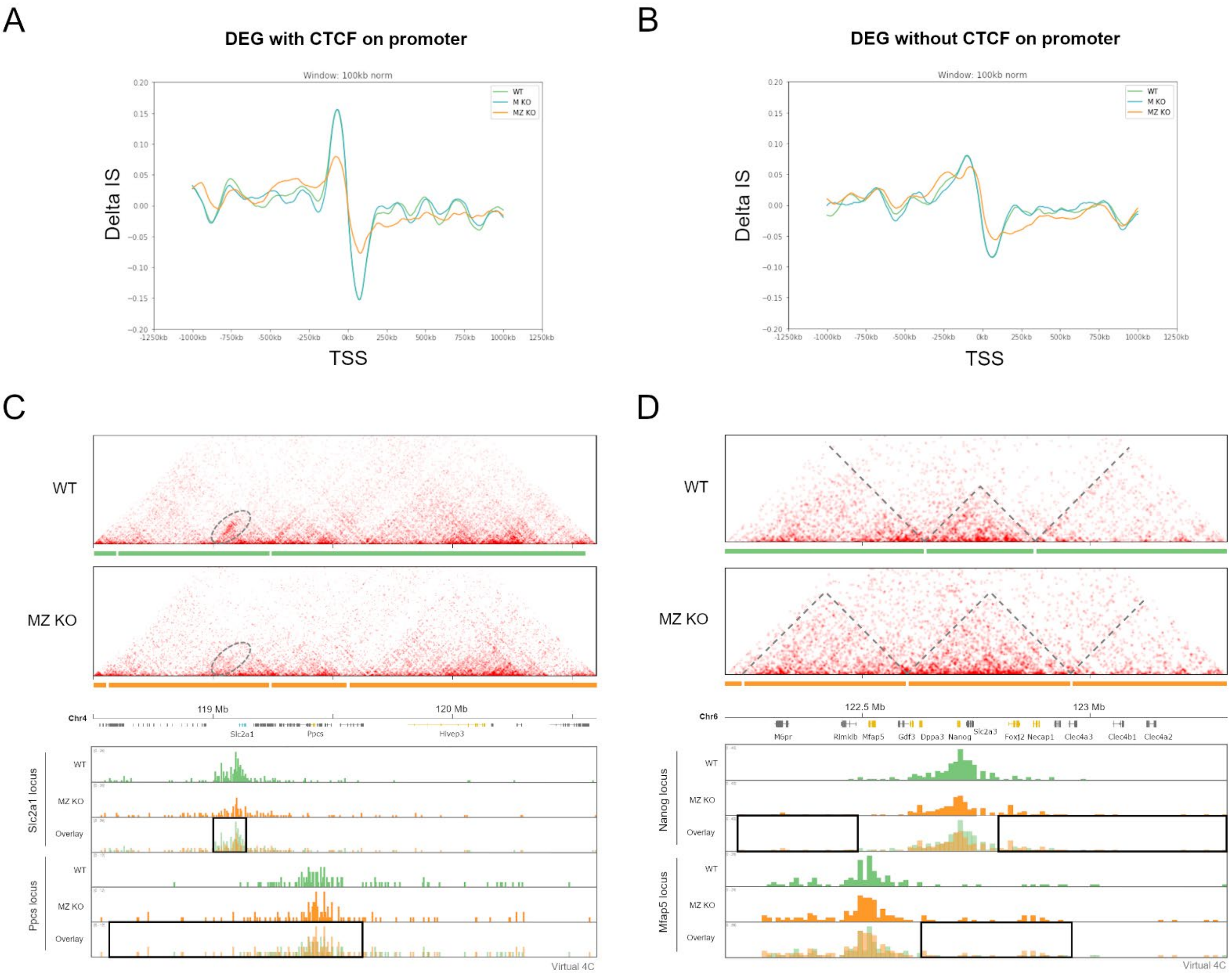


**Figure S4. Rewiring of contacts partially explains changes in transcription.** (***A***, ***B***) Changes in insulation score (Delta IS) around the transcriptional start site (TSS) of differentially expressed genes in MZ KO compared to control blastocysts with (***A***) or without (***B***) CTCF binding at their promoters, from WT (green), M KO (blue) and MZ KO (orange) datasets. Plots are centered at the TSS regions ±1000 kb. (***C***) Hi-C contact maps of the *Slc2a1* and *Ppcs* loci from WT and MZ KO blastocysts (upper panel). TAD domains in WT and MZ KO embryos are indicated below the matrices in green and orange, respectively. Only expressed genes are shown (downregulated genes in blue, upregulated genes in orange and non-differentially expressed genes in gray). Dashed ovals show regions where contacts are reduced in MZ KO embruos. The lower panel shows virtual 4C generated with Juicebox software for viewpoints at the promtoers of *Slc2a1* and *Ppcs* at 5 kb resolution. Gray boxes highlight changes in contacts. (***D***) Hi-C contact maps of the *Nanog* locus from WT and MZ KO blastocysts (upper panel). TAD domains in WT and MZ KO embryos are indicated below the matrices in green and orange, respectively. Only expressed genes are shown (upregulated genes in orange, non-differentially expressed genes in gray). The lower panel shows virtual 4C generated with Juicebox software for viewpoints at the *Nanog* and *Mfap5* promoters at 10 kb resolution. Gray boxes (lower panels) highlight a spread of the contacts in *Ctcf* mutant embryos and dashed lines (upper panel) highlight local changes in contacts.

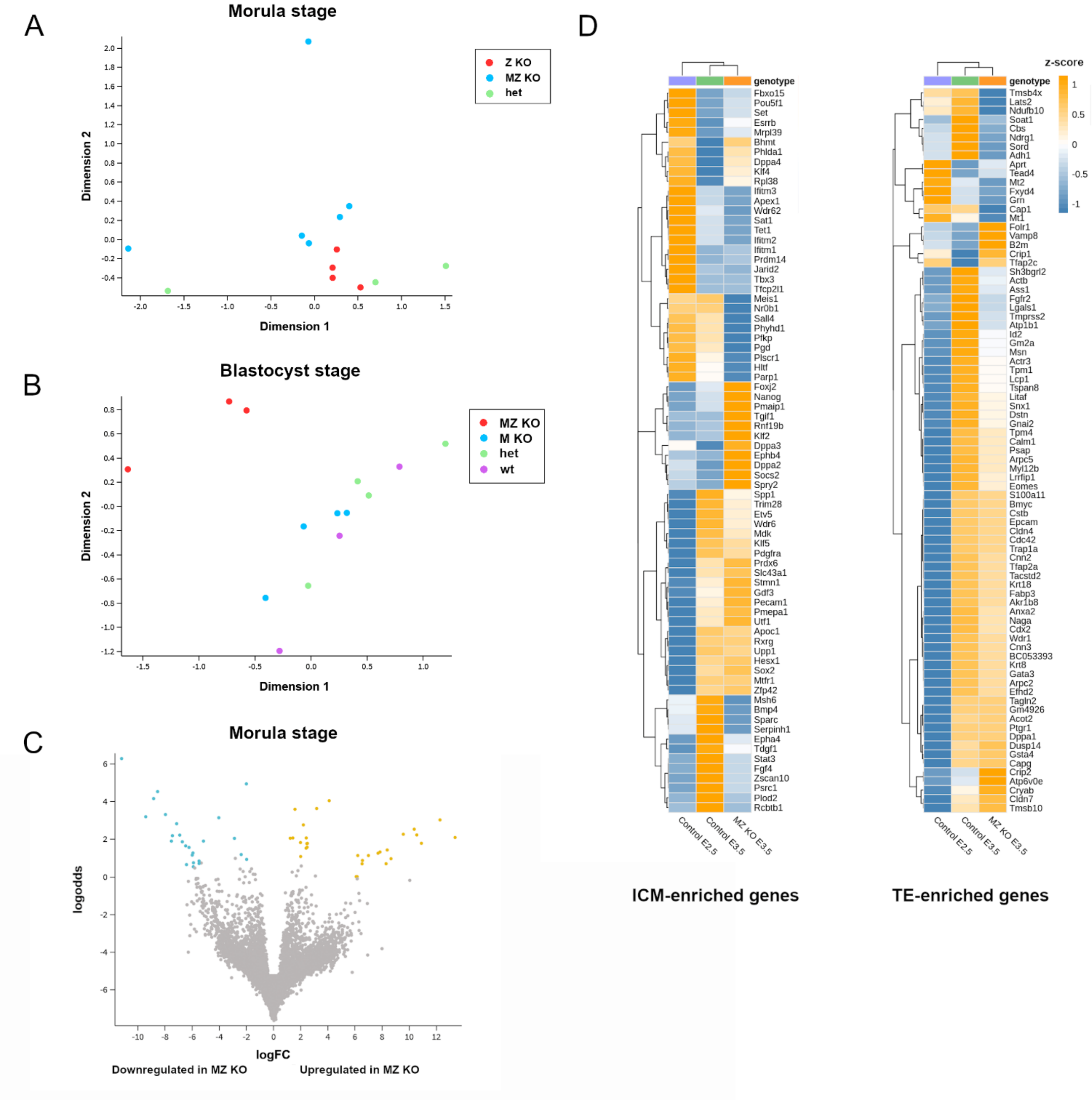


**Figure S5. Transcriptional changes in morula and blastocyst stage *Ctcf* mutant embryos.** (***A***) Multidimensional scaling -plot of RNAseq samples from single morula stage embryos of different genotypes. (***B***) Multidimensional scaling -plot of RNAseq samples from single blastocsyt stage embryos of different genotypes. (***C***) Volcano plot of differentially expressed genes between control ($Ctcf^{fl/-}$) and MZ KO single morulae. In blue, genes downregulated in MZ KO (adj. pvalue <0.05 and logFC <−1); in orange, genes upregulated in MZ KO (adj. pvalue <0.05 and logFC >1). Non-differentially expressed gens are shown in gray. (***D***) Heatmap showing the expression (z-score) of ICM- (left) and TE-enriched (right) genes from control morulae, control blastocysts and MZ KO blastocysts. Clustering of samples are shown on top.
